# Linguistic Rules of Syntax and Sharing in the Complex Songs of a Tropical *Passerine*

**DOI:** 10.1101/2025.07.31.666818

**Authors:** Suyash Sawant, Chiti Arvind, Viral Joshi, Indranil Dutta, V. V. Robin

**Affiliations:** Department of Biology, Indian Institute of Science Education and Research (IISER) Tirupati, Tirupati, AP 517619, India; Department of Wildlife Ecology and Conservation, University of Florida, Gainesville, FL 32603, USA; School of Natural Resources and Environment, University of Florida, Gainesville, FL 32603, USA; School of Languages and Linguistics, Jadavpur University, Kolkata, WB 700032, India

**Keywords:** Birdsongs, Cultural Evolution, Human Linguistics, Song Complexity, Song Sharing, Territoriality

## Abstract

1. Birdsong offers a powerful model to study the evolution of complex communication systems. While most song syntax and sharing research focuses on temperate species with fixed repertoires, we know far less about tropical songbirds with large, open-ended vocal systems.

2. We examined song structure and cultural transmission in the White-bellied Sholakili *Sholicola albiventris*, a tropical passerine endemic to the Shola Sky Islands of southern India. We studied song complexity and spectro-temporal variation using over 6,000 songs from 17 color-banded males across five years. We analyzed the song syntax using sequence-based metrics, including Levenshtein Distances and N-gram models.

3. Our results reveal high syntactic variability across short and long timescales, with stable note-type cores and structured expansions of combinatorial sequences. Higher-order N-gram turnover was rapid and strongly individual-specific, while broadly sharing the population-level note pool. Song sharing was significantly higher among neighbors than non-neighbors.

4. Using Generalized Additive Models, we explain how social and ecological factors, including neighbor count, territory area, repertoire size, N-gram usage, and the spectro-temporal structure of N-grams, determine consistency across years and sharing across individuals. This study highlights parallels between human linguistics and syntactic organization in the complex songs of a tropical passerine.

## 1. Introduction

Vocal communication is central to mediating social interactions, coordinating reproductive behaviors, and facilitating parental care across animal taxa (Janik and Slater 2000; Mariette 2019; Riebel et al. 2019). Songbirds (*Passerines*) are particularly recognized for their structural and functional complexity in vocalizations, making them a good model system for investigating the evolution of vocal communication. Songs in these species function as a reproductive strategy, enabling individuals to attract mates, compete with rivals, and establish individual identity (Kroodsma and Byers 1991; Brenowitz et al. 1997; Beecher 2017; Rose et al. 2022). These functions often rely on the recognition of song patterns, which allows individuals to assess one another and modulate their behavior through social interactions (Catchpole 1987; Leitão et al. 2022). In many species, song sharing is typically a result of vocal learning and social affiliation (Hughes et al. 1998; King and McGregor 2016; Sánchez and Mennill 2024), which implies knowledge of neighbours’ songs and contributes to maintaining territories. Song sharing, with group-level similarities and individual differences, creates a complex selective information landscape for vocal expression and cultural transmission (Slater 1986; Dhondt and Lambrechts 1992; Podos and Warren 2007).

Like human language, birdsong consists of sequences of discrete acoustic units: notes, syllables, or motifs arranged in non-random, often rule-based orders (Kuhl 2003; Kershenbaum et al. 2014; Williams 2021). This syntactic organization enables birds to generate a large variety of vocal outputs from a limited set of elements, allowing for both repetition and recombination, much like the phonemic and grammatical structures in human speech (Goldstein et al. 2003; Berwick et al. 2011; Samuels 2015; Kershenbaum et al. 2016). Although birdsong lacks semantic referentiality, it is often socially learned, context-dependent, and hierarchically organized, especially in species with complex social dynamics (Doupe and Kuhl 1999; Fishbein et al. 2020). These parallels have led to an expanding literature comparing avian vocal systems and human language in terms of learning mechanisms, sequence structure, and cultural evolution (Slater 1986; Griesser et al. 2018; Aamodt et al. 2020; Williams and Lachlan 2022).

However, most of what we know about syntactic rules in birdsong again comes from temperate species with relatively simple, closed-ended repertoires (Hughes et al. 1998) - birds that learn all their songs in a critical short window of development. It remains unclear whether similar structural rules apply to species with large, variable repertoires and open-ended learning, such as many tropical songbirds (Price and Yuan 2011; Araya-Salas and Wright 2013). These open-ended learners can acquire new vocal elements throughout their lives, resulting in diverse and context-dependent song structures, and allowing for individual identity, repertoire turnovers, and rapid cultural evolution (Cornez et al. 2017). This lifelong capacity for vocal modification parallels aspects of human language learning, where individuals can acquire new words, dialects, or even new languages well into adulthood (Hughes et al. 1998; Bruno et al. 2021). These similarities suggest that open-ended vocal learning may be key in supporting flexible, socially responsive communication systems across species. The ability to learn new vocal elements throughout life challenges the assumption of fixed syntactic rules, raising critical questions about social and ecological factors that shape the evolution of information landscapes in more dynamic and interactive environments.

In this study, we examine the structure and syntax of song sequences of a tropical passerine that is known to have highly variable songs - the White-bellied Sholakili *Sholicola albiventris*, endemic to the high-elevation Shola Sky Islands of the southern Western Ghats, India (Robin et al. 2017). This species is also thought to be an open-ended learner, making it a compelling system for studying variable vocal syntax and song sharing. We investigate how *S. albiventris* songs are constructed and modified over time, as well as their ecological significance. Specifically, we address (a) individual level variation within a year, including changes in consecutive songs, short- and long-term song plasticity; (b) individual level variation across years, focusing on annual repertoire turnover and long-term repertoire drift; and (c) across-individual patterns, including neighbor song sharing and individual vocal signatures. This framework allows us to evaluate vocal variation across multiple temporal and social scales.

We apply a multi-scale analytical framework to capture the complexity of vocal syntax and its variation, representing songs as structured sequences of discrete acoustic units (Kershenbaum et al. 2016). We fragment songs using individual note types and higher-order sequences to capture vocal organization at different levels, quantifying how birds construct, reuse, and recombine vocal elements across social and temporal scales. Our approach integrates methods developed initially in computational linguistics and information theory. We quantify the minimum number of insertions, deletions, or substitutions required to convert one sequence into another using Normalized Levenshtein Distance (NLD), a widely used metric in language processing (Garland et al. 2012; Kershenbaum and Garland 2015). We also use N-gram models, adapted from text and speech analysis, allowing us to examine patterns of vocal sequence structure and track the emergence, persistence, or loss of syntactic fragments over time (Somervuo and Harma 2004; Sidorov et al. 2014; Smith 2014). We integrated these tools with Generalized Additive Models (GAM) (Hastie and Tibshirani 1990) to assess how ecological or social factors shape vocal consistency.

## 2. Methods

### 2.1 Study Species and Site

This study examined songs of the White-bellied Sholakili (*Sholicola albiventris*), a monomorphic flycatcher endemic to the high-elevation Shola forests of the southern Western Ghats, India. We focused on *S. albiventris* males, who are highly vocal during the breeding season and are known for their complex and extensive repertoires used in territory defense and mate attraction (Purushotham and Robin 2016; Sawant et al. 2022). Our focal population is in and around a forest patch in Kodaikanal, adjacent to the Kodaikanal Wildlife Sanctuary (Palani Hills, Tamil Nadu, India), at an elevation of ∼2100 m. Our study spanned over a partially disturbed montane forest landscape with a mix of native tropical montane cloud forest trees and *Pinus* and *Eucalyptus* plantations close to a human settlement (Arasumani et al. 2019) (Fig. S1). We conducted the fieldwork during the breeding seasons, March-June, spanning five years, with observations from four years, 2019 and 2021 to 2023, bearing the field logistic constraints due to the COVID-19 pandemic in 2020 (Fig. S1).

### 2.2 Banding and Territory Mapping

In order to track and record individuals (Dunn and John Ralph 2004), we captured and color-banded adult birds using mist nets near known singing perches. Since the species exhibits no sexual dimorphism in plumage or size, we determined the sex of individuals using molecular techniques. Blood samples were collected via brachial venipuncture and stored in a Queen’s lysis buffer. We used established protocols (Robin et al. 2021) for DNA extraction using DNeasy blood and Tissue Kit (DNeasy Blood & Tissue Handbook 2023), molecular sexing using P2/P8 primers (Bantock et al. 2008), PCR amplification, and gel electrophoresis to distinguish male and female individuals, which allowed us to identify and focus analyses exclusively on males (Table S1).

We followed color-banded males throughout the breeding seasons to outline approximate individual territories. Using handheld GPS units and cellphones at each confirmed sighting, we aggregated locations to construct spatial outlines of breeding territories in QGIS v3.16.11 (QGIS Association). Later, we manually adjusted these boundaries based on more field observations, including natural boundaries and unoccupied gaps following (Suarez Sharma et al. 2024).

### 2.3 Acoustic Data Collection and Processing

We obtained high-quality recordings of focal individuals using either a Sennheiser ME66 shotgun microphone or a Wildtronics Pro-Mono parabolic microphone mounted to Zoom H6 or H4N digital recorders. We recorded *S. albiventris* songs from within 10 m of the focal individuals in the mornings, 5:00-12:00, and in the evenings, 15:00-19:00, under low wind and minimal anthropogenic noise conditions. We targeted approximately 150 songs per individual per year (described further below), resulting in a final dataset of >6000 songs from 17 males across the four years (Table S2). All recordings were single-channel and stored as uncompressed files (WAV files, 44.1 kHz sampling rate, 16-bit resolution).

All recordings were processed using Raven Pro v1.5 (K. Lisa Yang Center for Conservation Bioacoustics at the Cornell Lab of Ornithology), generating spectrograms with a 512-sample Hann window, 50% overlap, and a 512 DFT. Songs were manually segmented based on pauses greater than 2 seconds. Following established conventions in avian communication, SS annotated each vocal element (note) as a discrete acoustic unit separated by a visible temporal gap (supported by auditory confirmation), an approximate threshold of >0.02 seconds (Kershenbaum et al. 2016). For each note, we marked the start-end times and frequency bounds using visual inspection of the spectrogram.

### 2.4 Sampling Effort and Statistical Power

We determined the optimal number of songs to record per individual based on an initial rarefaction analysis of note-type accumulation curves (Fig. S2A). Using a pilot dataset of over 300 songs each for 12 individuals in a year, we observed that the number of unique note types plateaued between 120 and 180 songs per individual. Based on this asymptotic pattern, we adopted a standardized target of ∼150 songs per male per breeding season to capture most of each individual’s vocal repertoire while minimizing redundancy.

To ensure sufficient statistical power to detect biologically meaningful variation in song structure, we conducted sensitivity analyses using Analysis of Variance (ANOVA) in G*Power 3.1 (Faul et al. 2009). For temporal comparisons across four years within the same individual (n = 600 songs) and social comparisons across 12 individuals within a year (n = 1800 songs), our sampling design achieved 90% power to detect small effect sizes (*f* = 0.154 and *f* = 0.109, respectively). These results confirm that despite our species being an open-ended learner, with a high repertoire size, our dataset is sufficiently sensitive to detect fine-scale variation in vocal traits across both temporal and ecological dimensions (Fig. S2B-S2C).

### 2.5 Note Classification Pipeline

We implemented a semi-supervised classification pipeline using Spectrogram Cross-Correlation (SPCC) (Charif et al. 2010) to assign vocal elements to discrete note types and estimate repertoire diversity with a manually curated reference set of 1,000 notes, which served as the initial training dataset. The notes were annotated in Raven Pro v1.5 and exported, processed, and frequency bandpass filtered them using the *warbleR* package in R (Araya-Salas and Smith-Vidaurre 2017) to compute pairwise SPCC, with Pearson’s correlation used to quantify spectrogram similarity (Fig. S3).

We conducted the classification iteratively: each unclassified note was compared to reference notes until a correlation >0.97 assigned it a matching type. If the algorithm found no match, it accepted the highest-scoring match >0.8; otherwise, it treated the note as a new type, assigned a novel label, and added it to the training set. We selected these cut-offs based on manual validation of classified types of ∼1000 notes, cross-validated by two independent researchers (SS and CA). The algorithm added up to five replicates per newly assigned type to the training set to capture within-type variability (Fig. S3). The dataset was incrementally updated as new types emerged, followed by a manual review to ensure structural distinctiveness, collapse redundant classes, and assess classification accuracy through random spot-checks. In all, we manually assessed around 2% of the notes, and with ∼160 note types among 46,628 annotated notes, we felt this method of automation combined with manual verification was the only practical way to conduct this procedure in a scalable manner.

### 2.6 Quantifying Song Features

#### 2.6.1 Song Structure & Complexity Metrics

We used custom R scripts (Sawant 2025) to summarize the song-level spectro-temporal structure from extracted spectro-temporal parameters for each annotated note, including low, mean, and high frequencies, frequency bandwidth, and duration. We used both note- and song-level parameters to examine acoustic parameters across individuals and years (Fig. 1). We calculated a range of metrics capturing both within-song variation and across-song repertoire structure to quantify vocal complexity. Within each song, we measured note count (total number of notes), note pace (notes per second), note types (number of distinct note categories), note richness (proportion of note types), and note diversity using the Song Richness Index (SRI) (Sawant et al. 2021). We assessed note order using first-order Markov Entropy Rates (MER) (Kershenbaum 2014) to evaluate syntactic predictability and spectro-temporal variation using a Note Variability Index (NVI) (Sawant et al. 2022) based on spectro-temporal dissimilarity among notes (Fig. 1). To evaluate across-song repertoire structure, we estimated individual-level note repertoires using the Chao1 estimator in the vegan package (Kindt 2020) in R, which accounts for undetected note types. We also generated rarefaction curves to assess note-type accumulation across increasing song samples, allowing us to evaluate repertoire saturation and compare vocal richness for an individual within a year.

**Figure 1:**
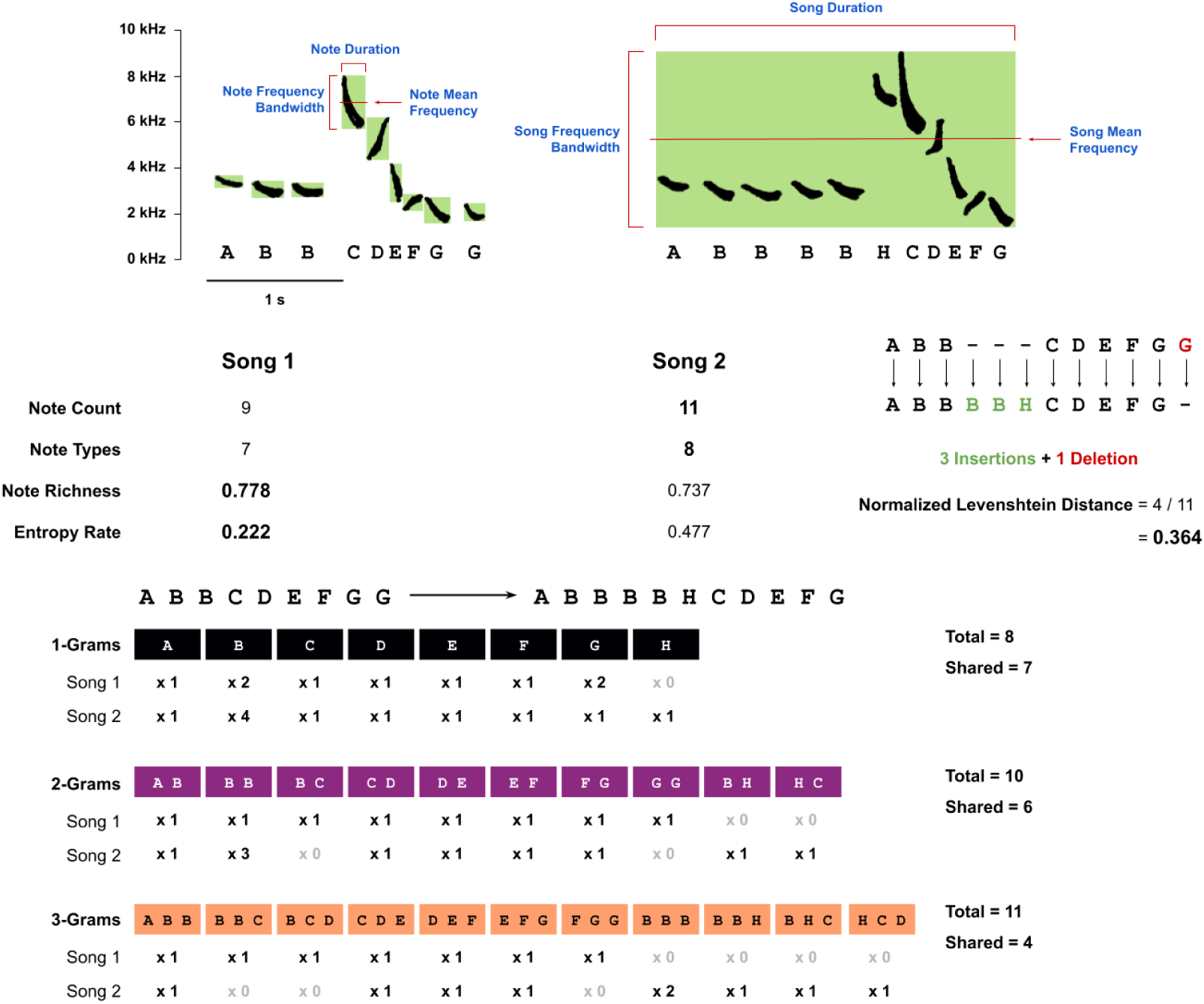
Comparison of song structure and syntax using two example song sequences. Spectrograms show two songs (Song 1 and Song 2) with labelled note identities and acoustic annotations at both the note and song levels. Key spectro-temporal parameters - duration, mean frequency, and frequency bandwidth are illustrated on the spectrograms. The summary metrics below each song quantify vocal complexity using specific complexity metrics. The Normalized Levenshtein Distance (NLD) measures syntactic dissimilarity between the sequences based on the number of edit operations required for alignment. The lower panel splits each song into its constituent N-grams:1 grams (individual notes), 2-grams (note pairs), and 3-grams (note triplets). Counts shown beneath each N-gram indicate their frequency within each song.

#### 2.6.2 Song Syntax Metrics

To quantify fine-scale structural differences between songs, we calculated Normalized Levenshtein Distance (NLD). This string-edit metric estimates the minimum number of insertions, deletions, or substitutions required to convert one sequence into another. We treated each song as a linear sequence of note-type names and computed pairwise NLD between songs produced by the same individual within a recording session, across sessions, and years (Fig. 1). These distances allowed us to quantify the degree of song-to-song variation and assess temporal patterns of vocal plasticity within bouts and breeding seasons. We compared consecutive and non-consecutive (random) song pairs to test whether syntactic changes were gradual or abrupt and whether divergence changed with time.

We extracted N-gram sequences from annotated notes to characterize higher-order syntactic structure, treating note types as discrete symbolic units. We focused on 1-gram (individual notes), 2-gram (adjacent note pairs), and 3-gram (adjacent note triplets) levels to capture increasingly complex patterns of syntax organization (Fig. 1). We compiled repertoires of N-gram types for each individual per year and calculated their diversity, frequency of use, and overlap across years. To assess repertoire turnover, we computed the proportion of N-grams retained, dropped, or newly introduced from one year to the next, providing insights into how vocal syntax evolves within individuals. To evaluate patterns of N-gram sharing across individuals, we measured pairwise overlap in N-gram repertoires. We also constructed cumulative N-gram accumulation curves across individuals to assess how syntactic diversity scales at the population level.

### 2.7 Modeling Song Sharing: Statistical Methods

To examine the ecological and acoustic drivers of syntactic structure in *S. albiventris* songs, we constructed two sets of Generalized Additive Models (GAMs) using the mgcv package in R.

#### 2.7.1 N-gram Consistency Across Years

The first model set focused on intra-individual consistency of N-gram usage across consecutive years. Each N-gram type was classified as either “consistent” (used in both years) or “not-consistent” (used in Year 1 - Y1 but not in Year 2 - Y2). We modeled the probability of each N-gram being classified as *consistent* or *learned* using the following predictors: (1) ΔTerritoryArea_Y1→Y2, i_: proportional change in territory area from Y1 to Y2 for *i*; (2) NeighborCount_Y1 ∪ Y2, i_: cumulative number of neighbors across the two years for individual *i*; (3) NgramOccurrence_Y1, i_: number of times N-gram *g* occurred in *i’s* songs in Y1; (4) NgramRichness_Y1, i_: total number of unique N-grams used by *i* in Y1; (5) NgramPosition_g_: mean position of N-gram *g* within songs; (6) NgramMeanFreq_g_: mean frequency (Hz) of N-gram *g*; (7) NgramFreqBand_g_: frequency bandwidth (Hz) of N-gram *g*; and (8) NgramDuration_g_: temporal duration (s) of N-gram *g*.

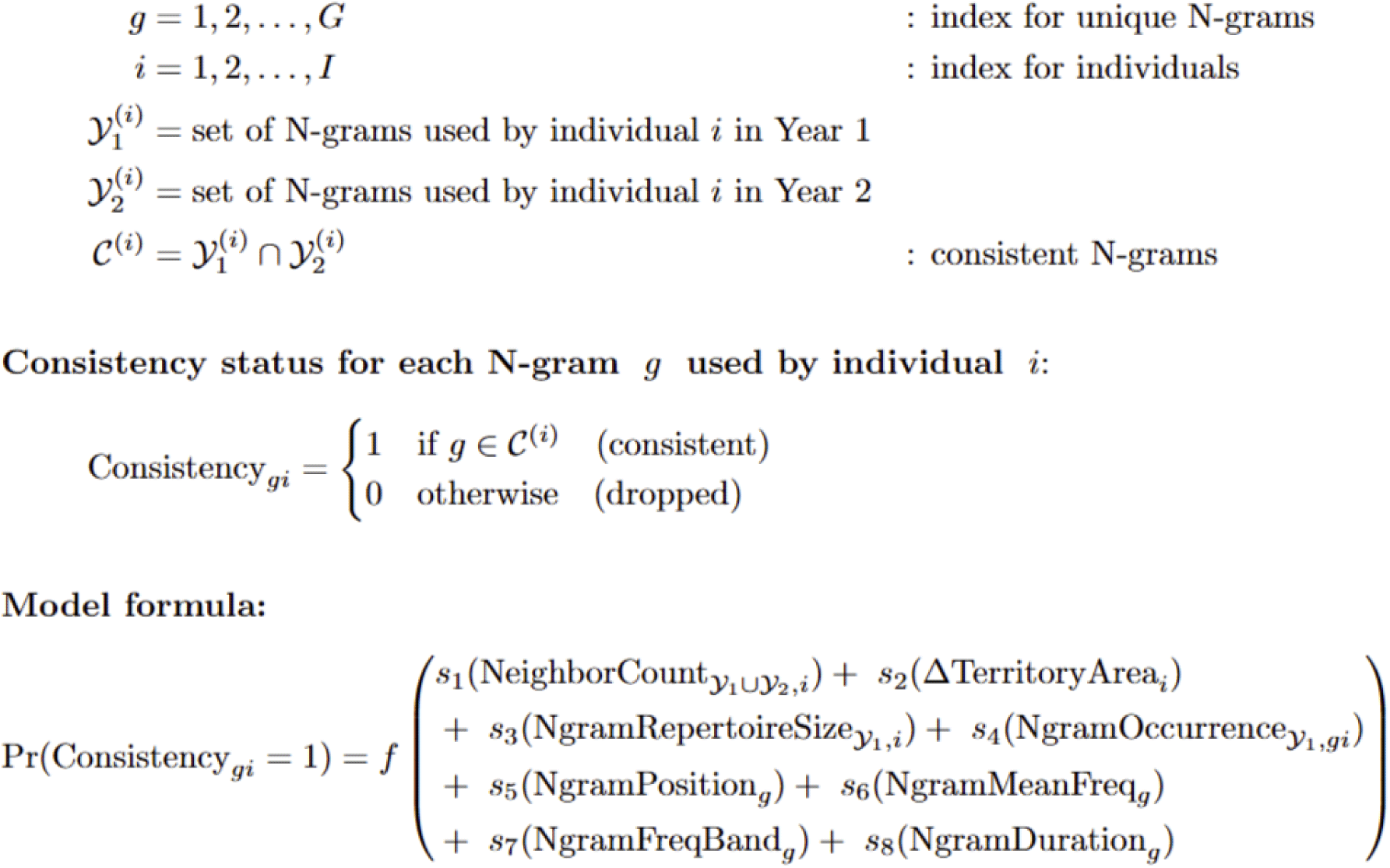

#### 2.7.2 N-gram Sharing Between Individuals

The second model set analyzed pairwise N-gram sharing within a year, where each N-gram was classified as “shared” (used by both individuals in a pair) or “not-shared” (used by only one of them). We modeled the probability of sharing between individuals *i*₁ and *i*₂ as a function of: (1) TerritoryDistance_i₁↔i₂_: distance (in meters) between territory centroids of the pair; (2) Shared NeighborCount_i₁ ∩ i₂_: number of shared neighbors between individuals *i*₁ and *i*₂; (3) max(NgramOccurrence_g, i₁_, NgramOccurrence_g, i₂_): maximum usage count of N-gram *g* between the pair; (4) NgramRichness_i₁ ∪ i₂_: total number of unique N-grams used by the pair; (5) NgramPosition_g_: mean position of N-gram *g* within songs; (6) NgramMeanFreq_g_: mean frequency (Hz) of N-gram *g*; (7) NgramFreqBand_g_: frequency bandwidth (Hz) of N-gram *g*; and (8) NgramDuration_g_: temporal duration (s) of N-gram *g*.

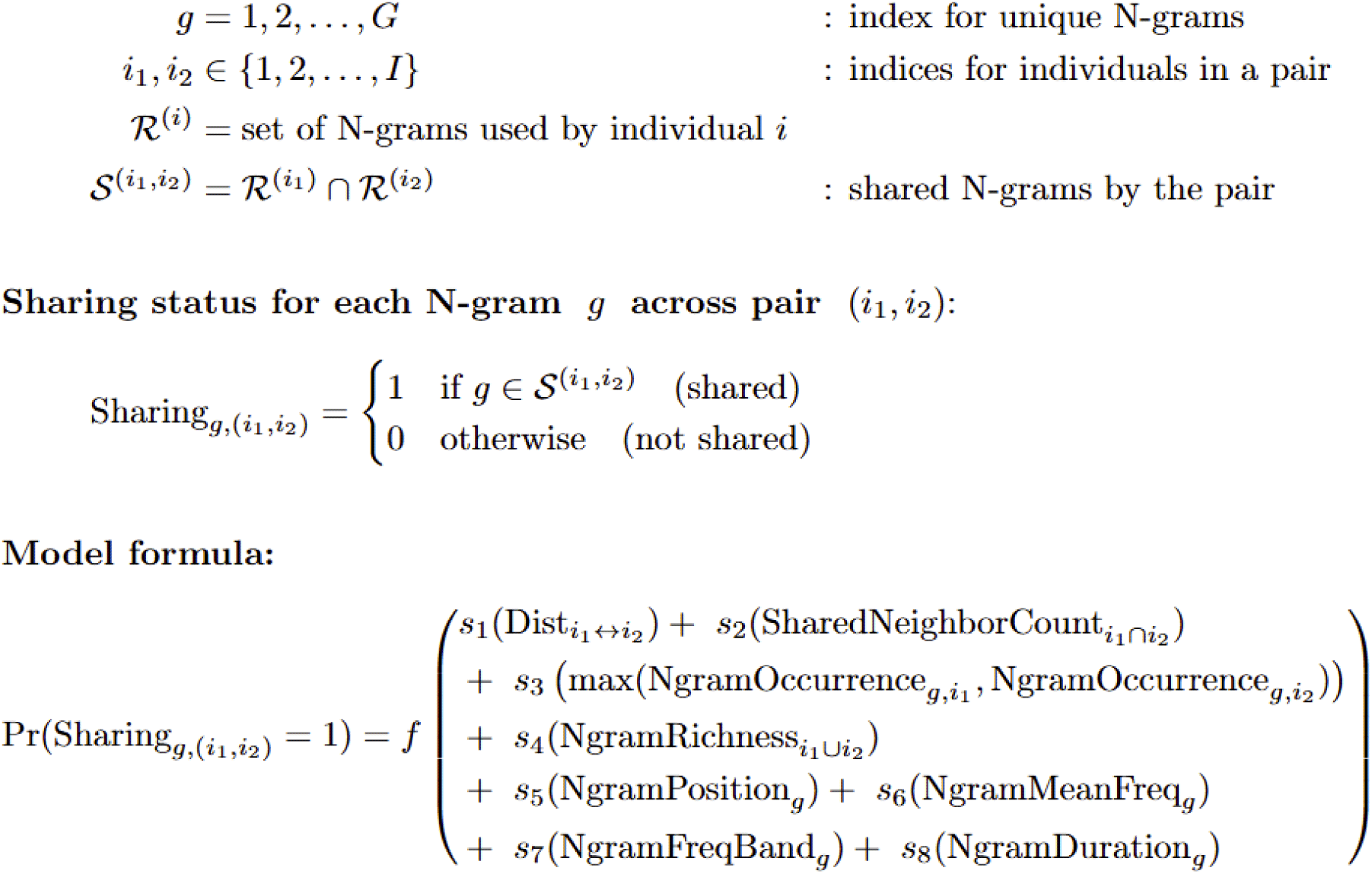

We modeled all predictors using penalized thin plate regression splines *s()*, allowing for flexible estimation of non-linear relationships. We ran all models separately for 1-gram, 2-gram, and 3-gram levels, and retained all predictors across models regardless of statistical significance to enable effect comparisons.

## 3. Results

This study analyzed over 6000 songs from 17 color-banded male *S. albiventris*, recorded over four breeding seasons (2019, 2021-2023). Using a combination of spectral, syntactic, and combinatorial metrics, we characterized vocal variation across three temporal scales within and across days, and across years. We examined spectro-temporal features such as mean and delta frequency and vocal complexity metrics such as note count, richness, and Markov Entropy Rate. In parallel, we quantified sequence-level consistency using Normalized Levenshtein Distances (NLDs) and tracked the growth of individual N-gram repertoires (unigrams, bigrams, trigrams) to assess structural plasticity over time. Finally, we evaluated social and spatial predictors of N-gram overlap among individuals to understand the extent and drivers of vocal sharing. These results offer a multi-layered perspective on the individual, social, and temporal dynamics shaping vocal communication in a montane songbird with highly variable songs.

### 3.1 Territorial Dynamics and Social Context

Despite the high site fidelity of many individuals (e.g., individual YYXX), we observed variation in territory size, locations, and neighboring individual interactions across four years (e.g., individual YOXX). For example, several individuals had turnover of neighboring males between years, and some territories declined in area. In contrast, we observed some new individuals (e.g., individual RUGG) dispersing into new areas, claiming empty spaces and/or pushing previously territory-holding individuals to introduce their territories. The long-term monitoring and banding allowed us to track persistent and newly recorded individuals in subsequent years (Fig. 2).

**Figure 2:**
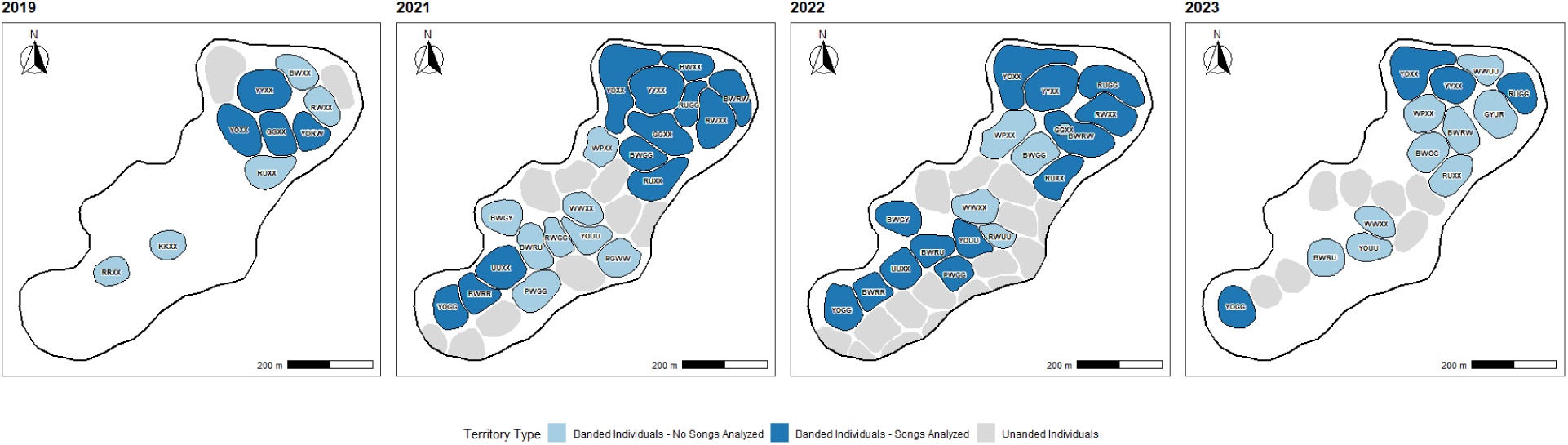
Annual (breeding season) territory maps of individually banded male *S. albiventris* across four breeding seasons (2019, 2021-2023) in the montane habitats of Kodaikanal, southern India. Polygons represent mapped male territories, with colors indicating banding and song analysis status: light blue = banded individuals, dark blue = banded individuals with song data analyzed that year, and gray = approximate territories of unbanded individuals. Territory boundaries are based on repeated resighting, spatial mapping, detecting song perches, and individual interactions, during the breeding season of the year.

### 3.2 Song Structure: Spectro-Temporal Features and Complexity

*S. albiventris* songs covered a broad frequency bandwidth of 3 to 9 kHz around a mean frequency between 4.5 and 5.5 kHz, with most individuals singing song-segments of 1.5 to 4 seconds duration. Despite this broad range, variation in these acoustic features was only partially structured by individual identity or year. Kruskal-Wallis tests revealed significant but moderate individual-level differences in mean frequency and bandwidth, suggesting that some males consistently favored higher or broader frequency ranges. Song duration also varied significantly across individuals, with some producing notably longer songs. Some individuals also showed significant differences in these spectro-temporal parameters across years (Fig. S4). The first two principal components resulted from the PCA of all song-level acoustic parameters, explained 87% of the variance, but did not indicate any distinct clustering by individual or year (Fig. S5). These results suggest spectro-temporal features alone do not encode individual identity or population dialect patterns.

Next, we examined song complexity metrics to assess vocal richness, diversity, and syntactic organization variation. These included note count, pace, repertoire size (note types and richness), diversity (Song Richness Index), spectro-temporal variation (Note Variability Index), and note order (Markov Entropy Rate). Most individuals showed significant differences in metrics, particularly in note count, note types, and song richness index. Some males consistently produced longer or more diverse songs, while others exhibited more repetitive or shorter sequences. Inter-annual variation was also evident in a few individuals, though overall patterns were more strongly structured by individual identity than by year (Fig. S6). A principal component analysis of all song complexity features revealed that the first two dimensions explained 82% of the variance, with PC1 capturing structural diversity, and PC2 reflecting note count and pace (Fig. S7). However, consistent with the spectro-temporal analysis, the PCA scores showed broad overlap among individuals and years, and did not reveal strong clustering by identity or time.

### 3.3 Song Syntax: Sequential Organization and Temporal Consistency

Despite a shared structural framework, *S. albiventris* songs exhibit striking internal diversity in their syntactic arrangements. Our analysis of normalized Levenshtein Distance (NLD) between song pairs sampled across three temporal scales within a day, across days within a season, and across years revealed consistently high values (typically NLD > 0.9; Fig. 3A), reflecting substantial syntactic plasticity. NLD values increased subtly with longer inter-song intervals beyond 4-5 minutes, suggesting that songs become progressively more distinct as temporal separation increases. Even within a single singing bout (generally a few minutes long), males rarely repeated similar song syntax (NLD < 0.5), and song pairs separated by more than a few minutes typically showed substantial structural divergence.

**Figure 3:**
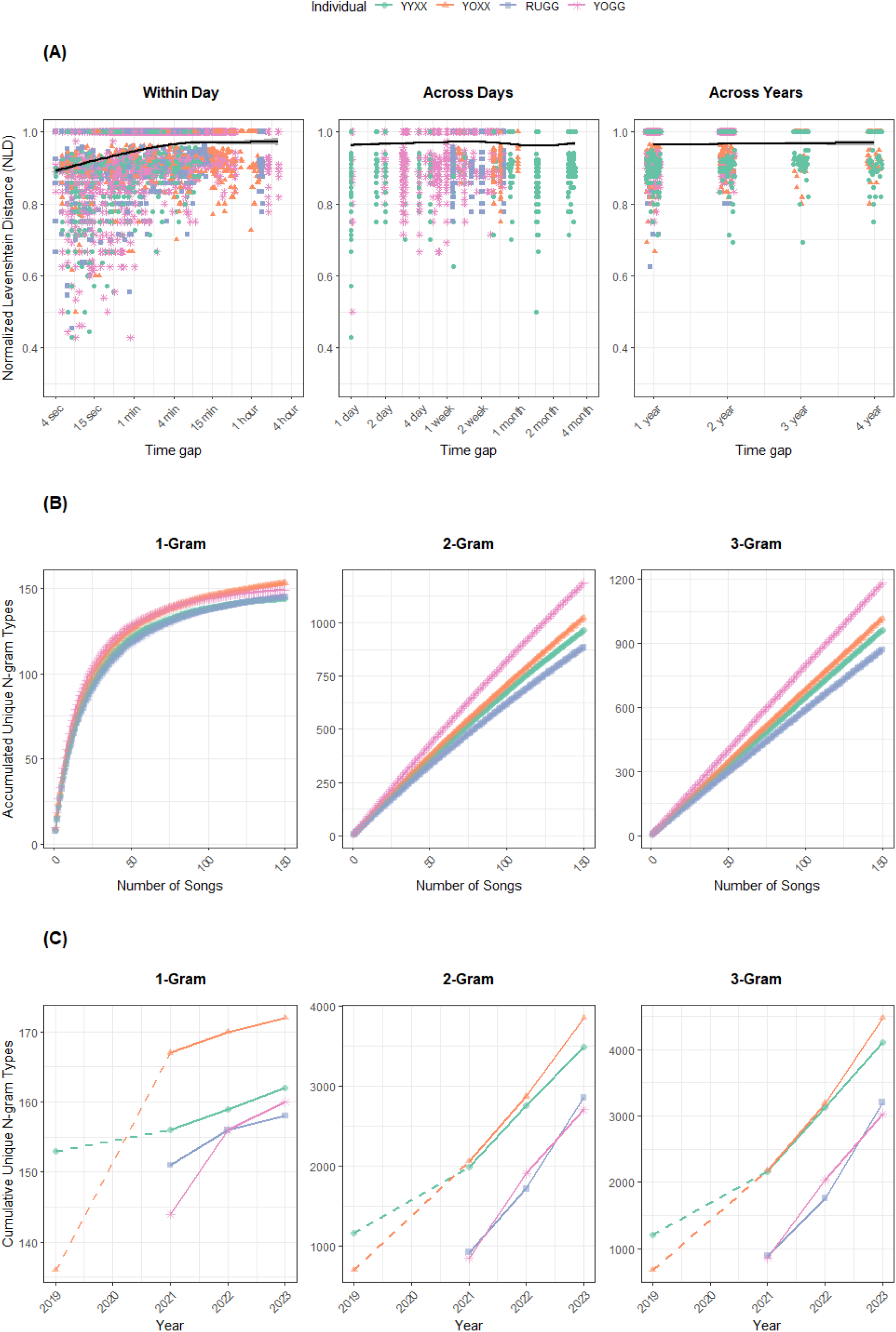
Song consistency and cumulative N-gram usage over time for four individuals. (A) Normalized Levenshtein Distance (NLD) between a random pair of songs as a function of temporal separation. Left: comparisons within the same day (log-scaled in seconds), middle: across days within the same year (log-scaled in days), right: across years (1-4 year gaps). Colored points indicate individual comparisons; black lines show smoothed GAM trends. (B) Repertoire accumulation curves for 1-gram, 2-gram, and 3-gram types in 2022. Each panel shows the number of unique N-grams accumulated as a function of the number of songs sampled. Curves represent individual birds, with ribbons indicating ±1 SD based on 100 random permutations. (C) Cumulative unique N-gram types across recording years. Solid lines connect consecutive years with available data; dashed lines represent gaps due to missing recordings in 2020. Each panel shows the cumulative number of distinct N-grams detected for each individual by year.

However, this high variation does not equate to disorder. Song syntax retained an underlying structure, particularly at the level of note types (1-grams). As we sampled more songs per individual, the number of unique 1-grams, 2-grams, and 3-grams steadily increased, although the rate and asymptote varied by N-gram level (Fig. 3B). Unigram repertoires saturated around 150 songs, suggesting a relatively stable note inventory. In contrast, the accumulation of bigrams and trigrams continued at a near-linear rate with increasing song count, indicating that novel combinations of notes emerge even within a single season.

By extending this analysis across years, we quantified the cumulative number of unique unigrams, bigrams, and trigrams produced over time (Fig. 3C). Unigram repertoires showed relatively shallow growth, with most note types reused across years and few novel notes added, indicating a relatively stable basic inventory. In contrast, the number of bigrams and trigrams increased rapidly each successive year, suggesting ongoing reorganization and innovation in how existing notes are combined. Even so, given the vast number of potential permutations, observed combinations represented only a small fraction of the possible syntactic space, implying that song syntax is not randomly assembled but constrained by individual-specific or culturally transmitted rules.

### 3.4 Song Sharing

We observed a strong effect of social factors on N-gram repertoire structure and sharing. Across both years analyzed (2021 - 12 individuals and 2022 - 13 individuals), cumulative N-gram richness increased with the number of individuals, but at varying rates depending on N-gram size (Fig. 4A). One-gram richness showed signs of saturation (possibly around 15 individuals), whereas 2-grams and 3-grams exhibited a near-linear accumulation, indicating higher combinatorial diversity. Overall richness was consistently higher in 2021 compared to 2022 across all N-gram sizes, suggesting temporal drift in repertoire complexity.

**Figure 4:**
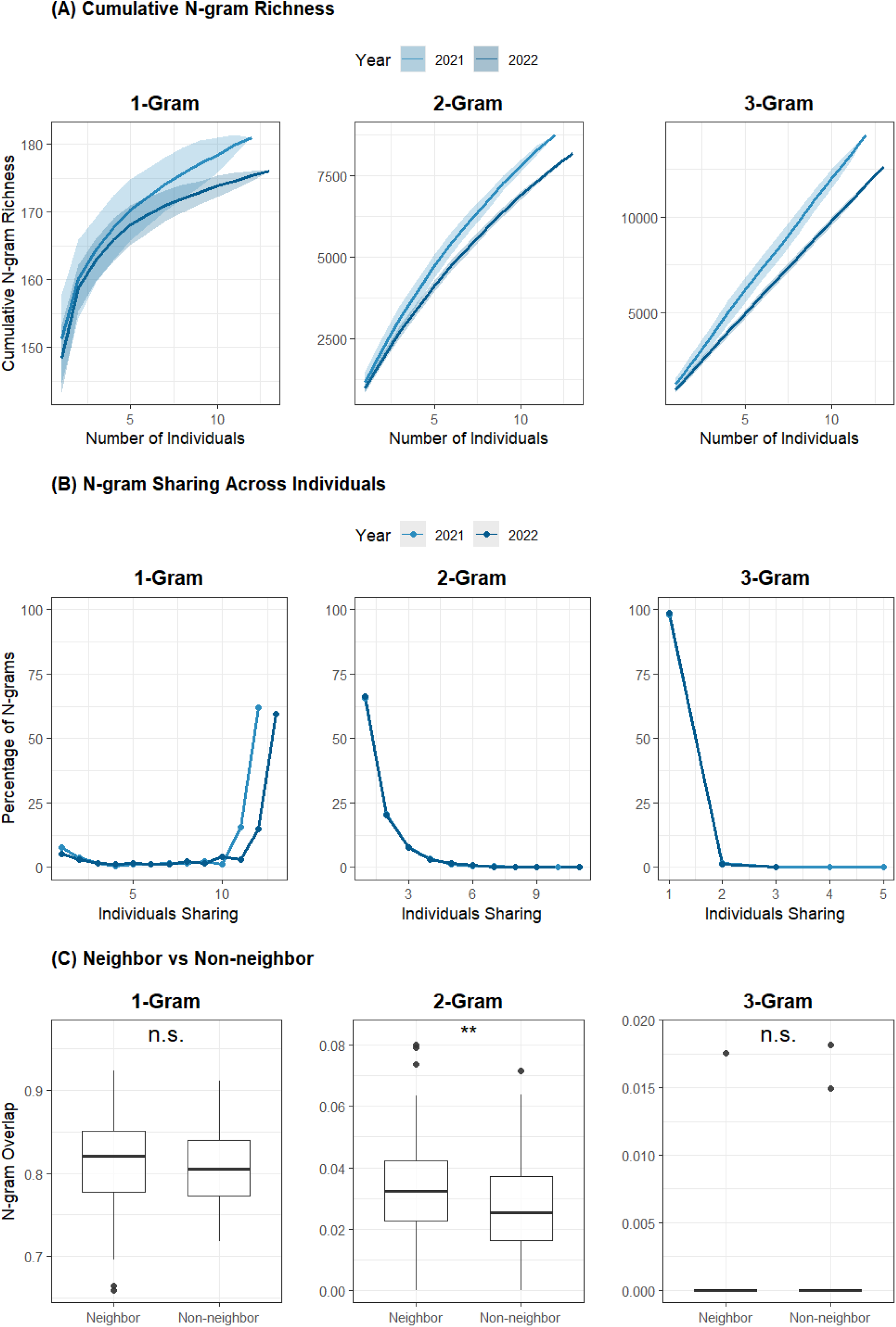
Social patterns of N-gram structure and sharing across individuals. (A) Cumulative N-gram richness as a function of the number of individuals, shown separately for 1-grams, 2-grams, and 3-grams. Lines represent mean richness across 100 random permutations, and shaded ribbons indicate standard deviation. (B) Distribution of N-grams by the number of individuals sharing them, shown as a percentage of total N-grams each year. Most 2- and 3-grams are individual-specific or rare. (C) Boxplots showing pairwise N-gram repertoire overlap between neighbors (territories ≤ 100 m apart) and non-neighbors, across N-gram sizes. Asterisks indicate Wilcoxon rank-sum test significance levels (* p < 0.05; ** p < 0.01; *** p < 0.001; n.s. = not significant). Year-specific comparisons are limited to 2021 and 2022, the two years with the most individuals recorded.

Despite this richness, most N-grams were rare and individual-specific. In both years, over 90% of 3-grams and over 75% of 2-grams were used by only one or two individuals (Fig. 4B). In contrast, a significant proportion of 1-grams was more widely shared, with all individuals sharing a subset of 1-grams. These patterns reflect a high degree of vocal individuality at higher-order N-gram levels, suggesting that complex syntactic sequences are not commonly shared, even among individuals in close proximity.

Consistent with this, N-gram repertoire overlap between neighboring individuals (territory centroids ≤ 200 m apart) was significantly higher than that between non-neighbors for 2-grams, indicating that shared short sequences may play a role in local social recognition or song matching (Fig. 4C). No significant difference was found for 1-grams and 3-grams, reinforcing that higher-order structures are more individualized and less influenced by spatial proximity.

### 3.5 Individual Signatures

We identified a set of individual-specific N-grams uniquely produced by only one focal male in a given year, highlighting the presence of individualized vocal signatures. For each focal individual (eg, YYXX, YOXX, RUGG, YOGG), we extracted the most frequently occurring 1-gram, 2-gram, and 3-gram types that met this criterion (Fig. 5). While all individuals exhibited distinct 2-gram and 3-gram signatures, 1-gram signatures were more limited, with some individuals (e.g., YYXX) lacking any consistent single-note signature in either year. The occurrence rates of these unique N-grams varied across individuals and years, but were relatively rare. The signature N-grams were structurally diverse, comprising short tonal elements and longer, frequency-modulated sequences. Many higher-order signatures included note combinations with distinct spectro-temporal features, suggesting that individual identity may be encoded not just in the presence of specific notes but also in the arrangement, duration, and acoustic structure of note sequences.

**Figure 5:**
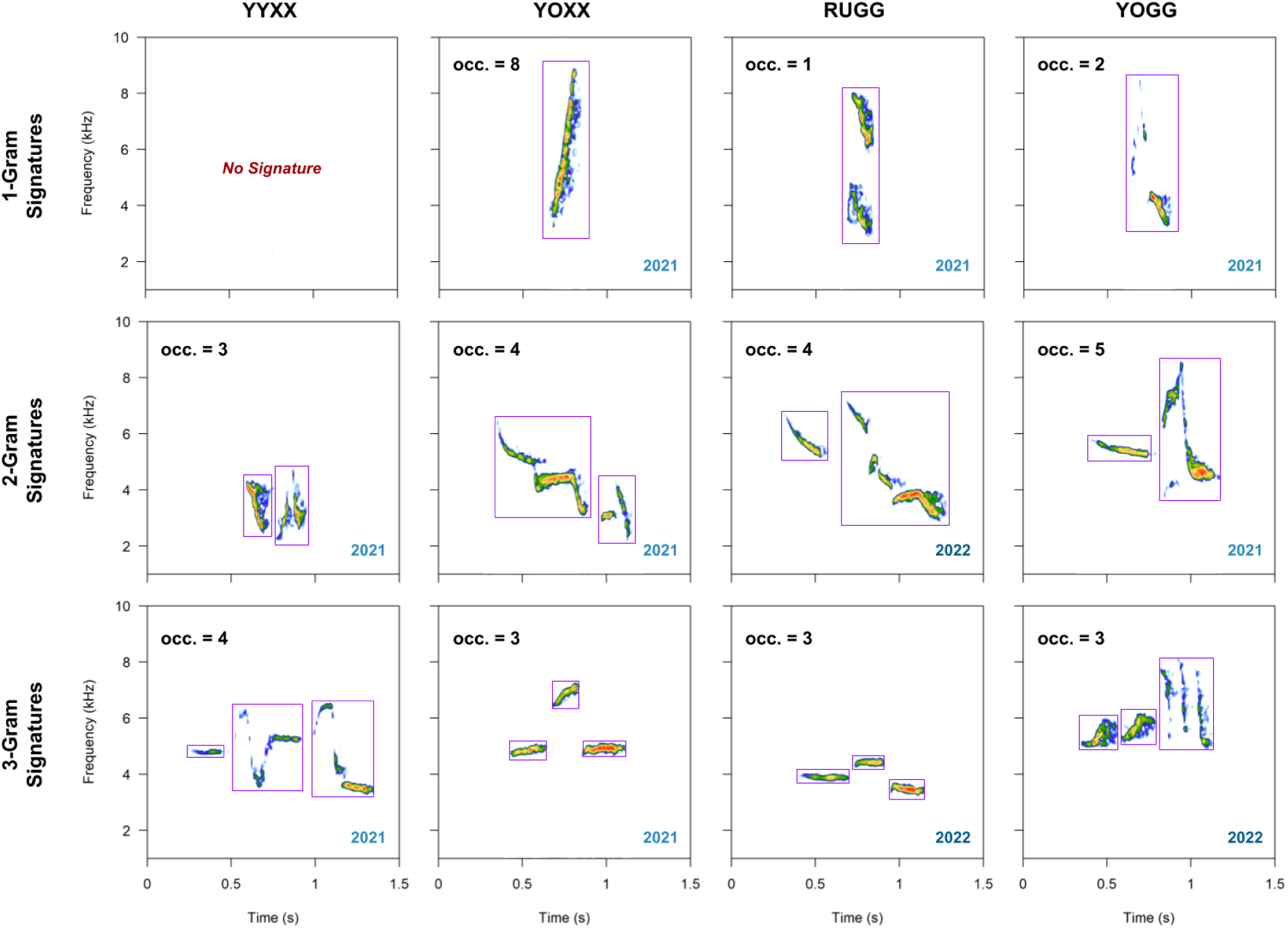
Individual-specific N-gram signatures in *S. albiventris* songs. Rows correspond to 1-gram, 2-gram, and 3-gram units; columns represent four focal individuals (YYXX, YOXX, RUGG, YOGG). Each panel displays a spectrogram of the most frequently occurring N-gram that was uniquely produced by that individual in a given year. The total number of occurrences across ∼150 songs is indicated at the top-left of each panel. The year of signature occurrence is shown at the bottom-right. Purple boxes outline individual notes comprising each N-gram.

### 3.6 Impact of Social and Ecological Factors on Song Consistency and Sharing

#### 3.6.1 Predictors of N-gram Consistency Across Years

Generalized additive models revealed that the probability of an N-gram being retained across years within individuals was significantly influenced by N-gram occurrence and repertoire richness in Year 1, as well as by acoustic features such as frequency bandwidth and mean frequency (Fig. 6A). N-grams with higher occurrence frequency in Year 1 were more likely to be retained, suggesting that frequently used elements form the core of an individual’s vocal repertoire. Repertoire richness showed a nonlinear relationship: individuals with intermediate-sized N-gram repertoires had the highest probabilities of retention. In contrast, ecological factors such as proportional change in territory area and total number of neighbors across years had weak or nonsignificant effects.

**Figure 6:**
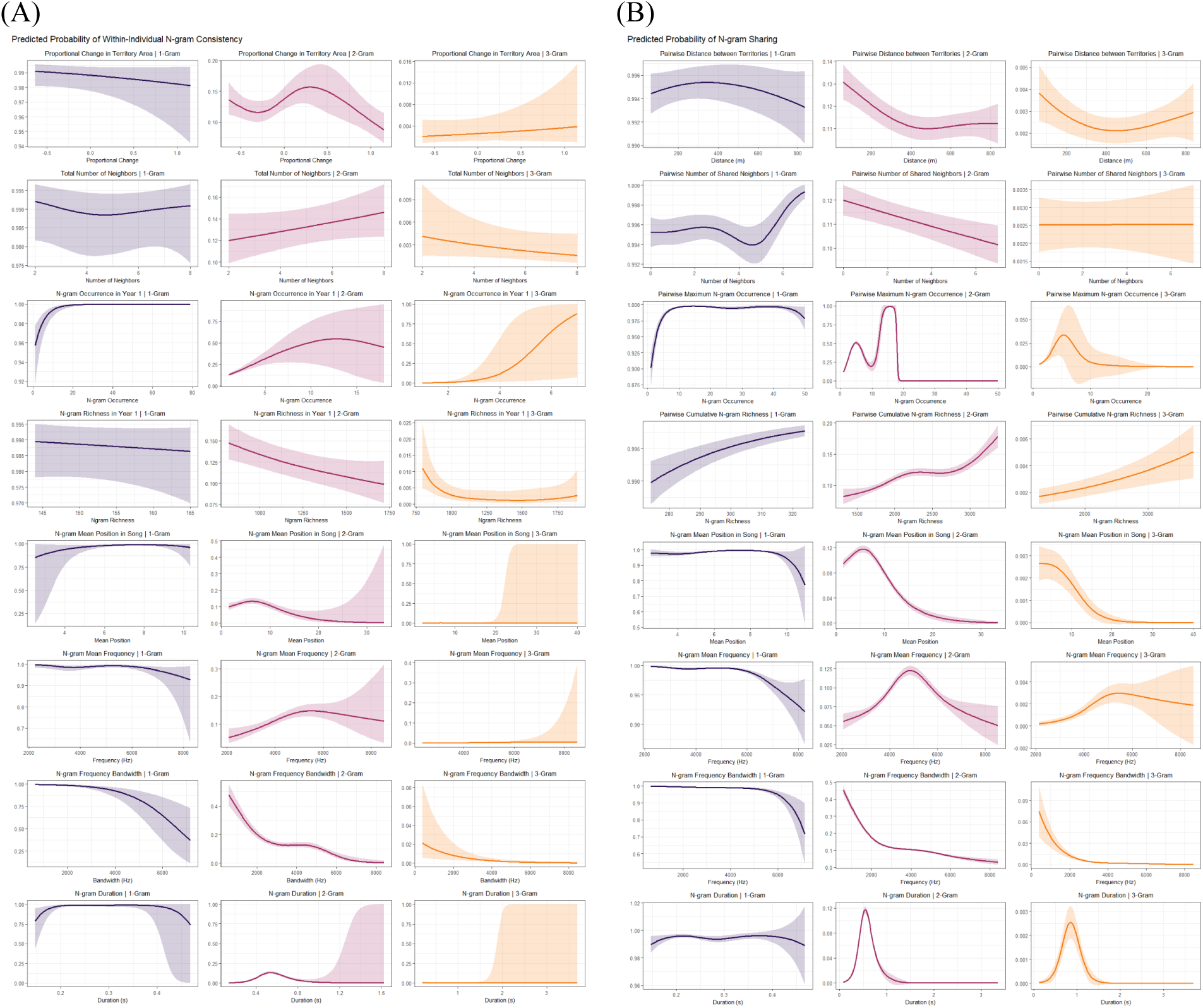
Results of generalized additive models (GAMs) examining ecological and acoustic predictors of N-gram usage in *Sholicola albiventris*. (A) Predicted probability of within-individual N-gram consistency across consecutive years. (B) Predicted probability of within-year N-gram sharing across individuals. Each column represents a different N-gram size (1-Gram, 2-Gram, 3-Gram), and each row shows the effect of a specific predictor on the probability of consistency or sharing. Rows 1-4 differ by model type: In (A) Consistency models, the predictors are: Proportional change in the territory area from Year 1 to Year 2, Total Number of neighboring individuals across two years, N-gram occurrence frequency in Year 1, and N-gram repertoire size in Year 1. In (B) Sharing models, the predictors are: Pairwise distance between territories, Number of shared neighboring individuals, Maximum N-gram occurrence between the pair, and Cumulative N-gram repertoire size across the pair. Rows 5-8 (shared across both model types) show effects of acoustic features: Mean position of the N-gram in the song, Mean frequency of the N-gram, Frequency bandwidth, and Duration of the N-gram. Lines show GAM-predicted probabilities, with shaded ribbons representing 95% confidence intervals. Line colors correspond to N-gram size: 1-Gram (purple), 2-Gram (pink), 3-Gram (orange).

#### 3.6.2 Predictors of N-gram Sharing Within Years

Within-year sharing of N-grams across individuals was primarily structured by spatial proximity and social connectivity (Fig. 6B). N-gram sharing decreased with increasing distance between territories, and the sharing of notes (1-grams) increased with the number of shared neighbors, indicating localized convergence driven by social exposure. Larger combined repertoire sizes were positively associated with sharing, suggesting that individuals with larger repertoires are more likely to have shared N-grams.

#### 3.6.3 Syntactic and Structural Constraints Across Both Models

The position of N-grams is critical in sharing, as N-grams occurring in the initial part of the song are likely to be shared by individuals. The spectro-temporal structure of N-grams influenced consistency and sharing patterns, with comparable trends across model sets. Frequency bandwidth was one of the strongest predictors: N-grams with narrower frequency ranges were more likely to be retained across years and shared across individuals, suggesting reduced complexity may favor stability and convergence. Mean frequency also showed moderate nonlinear effects, while duration had a weaker, inconsistent influence (Fig. 6).

## 4. Discussion

### 4.1 Complexity beyond Spectro-Temporal Features

Traditional acoustic features and complexity metrics quantify only partial aspects of song syntax diversity in *S. albiventris*. While spectro-temporal features like frequency bounds and song duration are helpful descriptors, they fail to capture the organizational intricacies that characterize the elaborate vocal repertoire of this species. Single-value indices such as note pace, note richness, Song Richness Index (SRI), Markov Entropy Rate (MER), and Note Variability Index (NVI) are informative for broad comparisons but are limited in scope. These metrics reduce complex vocal syntax into simplified summaries and overlook the internal structure and sequential transitions. In species with open-ended repertoires and high syntactic variation, such as *S. albiventris*, communicative complexity lies not just in how many elements exist, but in how they are combined and reused across time and social contexts (Kershenbaum et al. 2014). This study emphasizes the need to move beyond traditional acoustic parameters and summary indices and adopt sequence-based, syntactic approaches to understand the intricacies of vocal communication in complex song systems.

### 4.2 Balancing Vocal Innovation and Identity

*S. albiventris* displays a dynamic balance between vocal stability and innovation. Males reliably reused core note types across years, suggesting stable vocal “signatures,” yet higher-order N-gram structures showed ongoing turnover, reflecting active innovation within repertoires. This dual pattern suggests an evolutionary trade-off between the need for individual recognizability and the flexibility to diversify signals in response to shifting social or ecological pressures (Briefer et al. 2008; Smith-Vidaurre et al. 2020). The inter-annual variation and syntactic plasticity observed here are typical of open-ended learners capable of modifying song beyond early development (Cornez et al. 2017). This structured plasticity shows the potential for using birdsongs as a model to study how communication systems evolve under competing pressures for stability and novelty.

### 4.3 Social Context as a Driver of Vocal Learning

Vocal learning is prominently shaped by intrinsic developmental programs and the social environment in which individuals acquire and use their songs. We found that males with more territorial neighbors tended to exhibit higher N-gram richness and greater syntactic flexibility, suggesting that vocal innovation is partially influenced by local social context. Territory overlap, proximity to other males, and opportunities for repeated vocal interactions may encourage learning and modifying vocal sequences. Our results suggest that *S. albiventris* males may navigate these demands by combining shared population-level syntax with personalized vocal signatures, such as those in European Wrens (Catchpole and Rowell 1993), Skylarks (Briefer et al. 2012), Great Tits (Rivera-Gutierrez et al. 2010), and Splendid Sunbird (Payne 1978) through repeated interaction with their acoustic neighborhood. This socially responsive vocal behavior is not unique, as social context plays a key role in shaping vocal learning trajectories, from dialect formation in temperate sparrows (Hensel et al. 2022) to vocal convergence in parrots (Wright and Dahlin 2017) and cetaceans (Janik 2014). Rather than being fixed endpoints of early-life learning, repertoires may remain open to change throughout adulthood, especially in socially flexible, open-ended learner species (Cornez et al. 2017).

### 4.4 Cultural Transmission and Formation of Vocal Dialects

Patterns of N-gram sharing among *S. albiventris* males revealed a core set of syntactic elements widely used within a given year, indicating a shared cultural pool. These common N-grams, sustained through social learning and horizontal transmission among neighboring individuals, form the basis of a population-level dialect. Similar dynamics have been observed in the Yellow-naped Amazon (*Amazona auropalliata*), where consistent variation in contact calls coexists with the long-term stability of vocal dialects across populations (Wright and Wilkinson 2001; Wright et al. 2008). At the same time, individual *S. albiventris* exhibited distinct N-gram patterns, creating a layered system of shared and individual-specific note sequences. In many passerines, such cultural variation is shaped by geographic and habitat barriers (Laiolo and Tella 2005; Ortiz-Ramírez et al. 2016; Camacho-Alpízar et al. 2018), short dispersal distances (Searcy et al. 2002; Graham et al. 2017), and constrained social networks (Fernandez et al. 2017; Williams 2021), which can promote the emergence of localized dialects across fragmented landscapes. These findings highlight the potential of birdsong as a model for studying how geographic isolation, dispersal constraints, and habitat connectivity drive the evolution and divergence of cultural traits.

### 4.5 Parallels Between Birdsong and Human Language

The syntactic structure of *S. albiventris* songs revealed striking similarities with the organization of human languages. In human speech, phonemes and words serve as symbolic units flexibly combined into larger sequences, and similarly, individual notes and N-grams in birdsong act as compositional building blocks (Doupe and Kuhl 1999; Kershenbaum et al. 2016). Our analyses showed that these sequences are neither random nor rigid but exhibit structured transitions and recurrent patterns, reflecting principles akin to compositional syntax in language (Berwick et al. 2011). *S. albiventris* syntax balances regularity and flexibility, suggesting a rule-governed but adaptable communication system similar to human languages. Vocal innovation, social learning, and inter-annual turnover in repertoires parallel key mechanisms of human cultural transmission that make it a valuable model for understanding the origins and dynamics of complex communication systems.

### 4.6 Analyzing Complex Song Syntax: Challenges and Opportunities

Quantifying syntactic structure in birdsong introduces significant analytical challenges, particularly in species like *S. albiventris* that possess large, variable repertoires with flexible note ordering (Kershenbaum et al. 2014; Sawant et al. 2022). Traditional methods often rely on fixed or stereotyped syntactic structures and may fail to capture the underlying organization in open-ended learners. While N-gram models are widely used to detect recurring syllables (Sasahara et al. 2012; Smith 2014), they require careful implementation to distinguish meaningful syntactic patterns from chance co-occurrences, particularly in systems with high sequence variability. The huge combinatorial space of possible note arrangements can also lead to data sparsity, especially at higher N-gram levels. Moreover, classification accuracy in note labeling can compound this challenge: for example, with 80% precision at the single-note level, the accuracy of a 2-gram drops to ∼64%, and declines exponentially for longer sequences. These factors underscore the need for analytical tools that are both sensitive to latent structure and robust to observational noise. Despite these difficulties, our study shows we can detect robust patterns of N-gram reuse and structured innovation even in open-ended learners with high variability.

### 4.7 Long-term Monitoring of Tropical Birds and Future Directions

Long-term monitoring is essential in tropical systems, where extended breeding seasons (Renthlei et al. 2022), high adult survival (Karr et al. 1990), and complex social dynamics (Buskirk 1976) place unique pressures on vocal communication. Our multi-year study of *S. albiventris* demonstrates the value of sustained, individual-level acoustic monitoring for uncovering syntactic variation, vocal plasticity, and social learning. Crucially, such insights depend on the ability to track known individuals across years, highlighting the irreplaceable role of color-banding in studies of vocal communication and cultural transmission. We may fully understand the key processes, such as repertoire turnover, the maintenance of individual vocal signatures, and the development of population-level dialects, only through extended observations of individually marked birds. Unfortunately, recent policy changes have made obtaining permissions for color-banding endemic birds in India increasingly difficult, potentially undermining future efforts to study long-term vocal communication (Shanker et al. 2023). As habitat fragmentation and degradation intensify across tropical montane forests (Laurance et al. 2011), culturally transmitted traits essential for mate attraction, territoriality, and social cohesion may become increasingly fragile. Long-term, individual-based studies remain vital for understanding and conserving the cultural dimensions of avian biodiversity.

## Declaration of AI usage

During the analytical process for the manuscript, we used ChatGPT-4o to simplify and clean our R scripts. We analyzed all the data in the R environment, and SS and CA thoroughly reviewed the scripts before finalizing the workflow. We used Grammarly to improve the readability and clarity, and Paperpile to manage citations, but we did not use Generative AI tools to write this manuscript. So we take full responsibility for the text presented here.

## Author Contribution Statement

**Suyash Sawant:** Conceptualization, Data curation, Formal analysis, Investigation, Methodology, Software, Validation, Visualization, Writing - original draft, and Writing - review & editing. **Chiti Arvind:** Conceptualization, Data curation, Formal analysis, Investigation, Methodology, Software, Supervision, Validation, & Writing - review and editing. **Viral Joshi:** Conceptualization, Investigation, Methodology, Validation, & Writing - review and editing. **Indranil Dutta:** Methodology and Writing - review & editing. **Robin V. V.:** Conceptualization, Funding acquisition, Methodology, Project administration, Resources, Supervision, Writing - original draft, and Writing - review & editing.

## Acknowledgement

We thank Harikrishnan C.P., Ashwin Warudkar, Vinay K.L., Amrutha Rajan, Archita Sharma, Naman Goyal, Anuja Kanase, and Susmit Bansode for their field and lab work assistance; Yogeshwari and Kamraj for assistance in the field. We are grateful to Ramana Athreya, Jobin Varughese, Nandini Rajamani, Michael Pitzrick, Russ Charif, Laurel Symes, Guha Dharmarajan, and Kathryn Sieving for valuable discussions and feedback on study design, analytical approaches, and broader conceptual framing. We also thank the members of the Bird Ecology Lab at the Indian Institute of Science Education and Research (IISER) Tirupati, for their thoughtful inputs.

We sincerely appreciate the Tamil Nadu Forest Department and the Kodaikanal Forest Division for granting the permissions (permit no. WL5(A)/43781/2017) necessary to conduct fieldwork. We thank Kodaikanal International School (KIS) and the School’s Center for Environment and Humanity, particularly Corey Stixrud, Ashwin Fernandes, and Iti Maloney, for providing access to the KIS-IISER field station and additional logistical support. We also thank Ian Lockwood, Ramesh Daga, Billy and Arun Kolhatkar, and the Palani Hills Conservation Council for generously granting access to their lands during data collection.

## Supplementary Material

**Figure S1:**
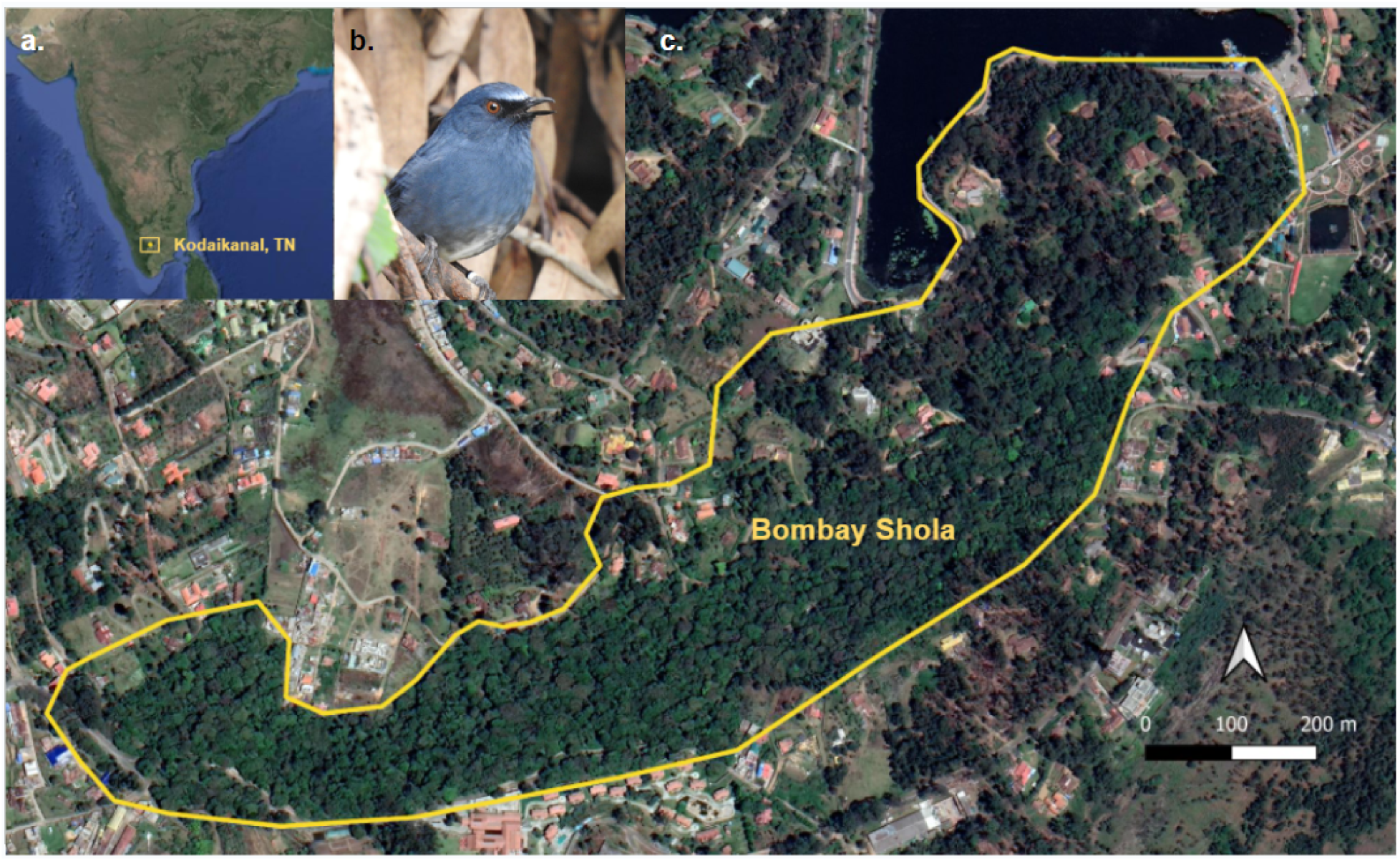
(a) Location of Kodaikanal, Tamil Nadu (TN), in the southern Western Ghats of India. (b) Adult male *Sholicola albiventris* (White-bellied Sholakili), the focal species individual with Black-White (BWXX) color-band. (c) Satellite view of the primary study site, Bombay Shola, a mid-elevation montane evergreen forest fragment (∼40 ha) adjacent to Kodaikanal town. The yellow polygon marks the forest boundary used for sampling and territory mapping.

**Figure S2:**
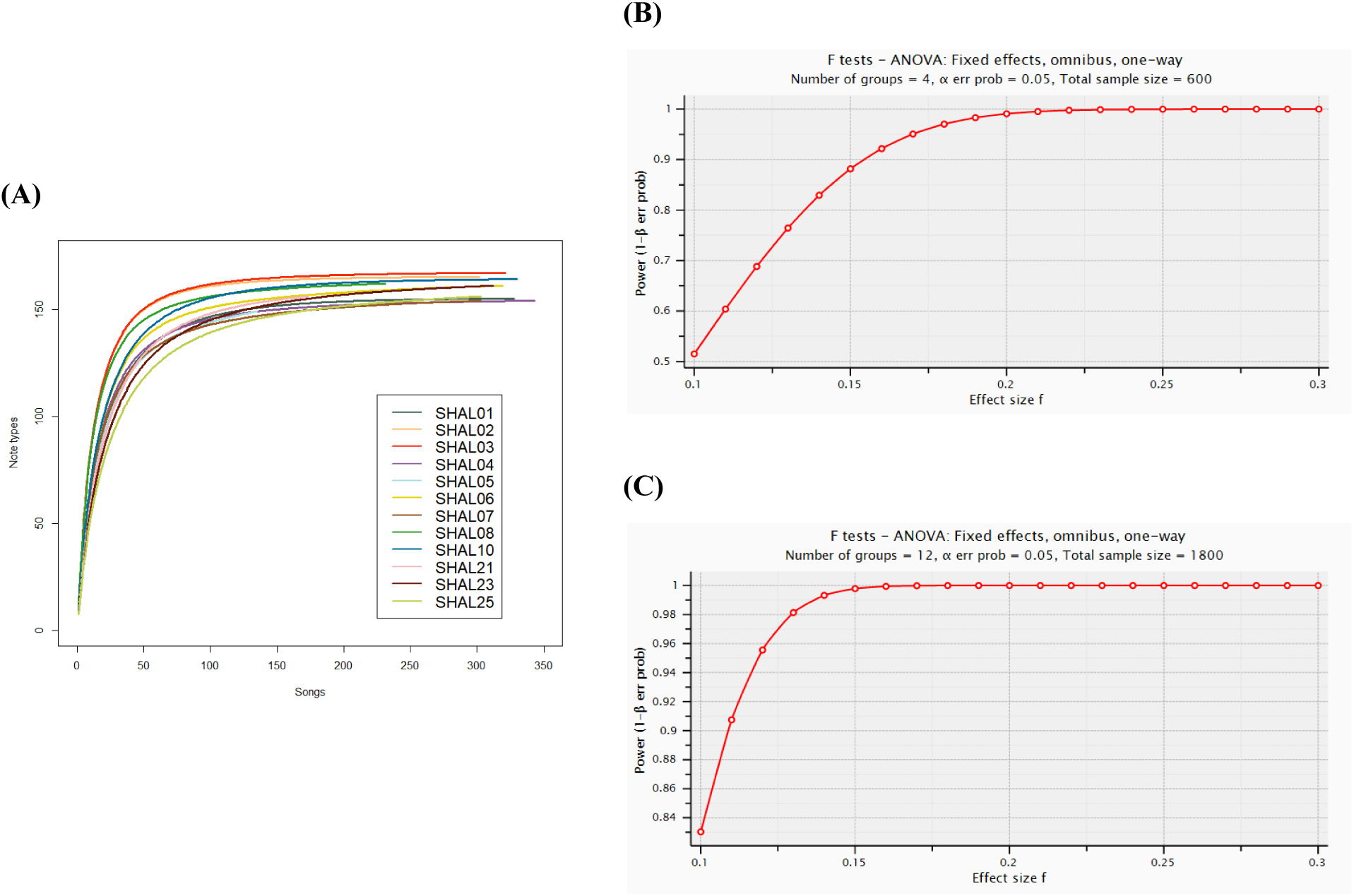
(A) Rarefaction curves showing note-type accumulation across songs for 12 males, indicating saturation between 120-180 songs per individual. (B) Power analysis for detecting within-individual variation across four years (n = 600 songs). (C) Power analysis for detecting among-individual variation within a year (n = 1800 songs). Power analyses were conducted using one-way ANOVA in G*Power 3.1, showing ≥90% power to detect small effect sizes.

**Figure S3:**
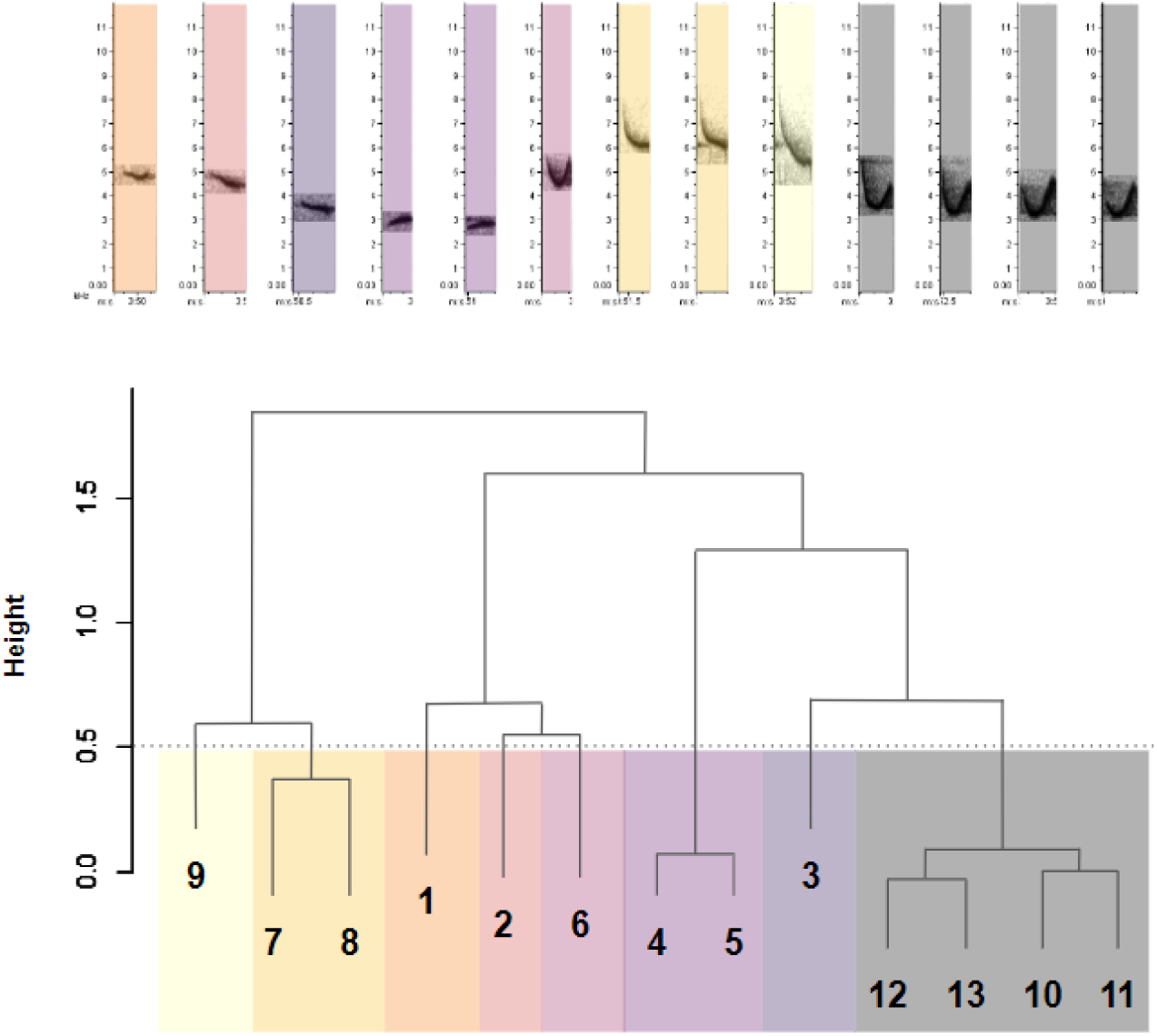
Representative spectrograms (top) and hierarchical clustering dendrogram (bottom) illustrating the classification of note types into 13 distinct acoustic categories based on similarity in acoustic features. Clustering was performed using hierarchical clustering using spectrogram cross-correlation. Colors correspond to the final note clusters used in downstream analyses.

**Figure S4:**
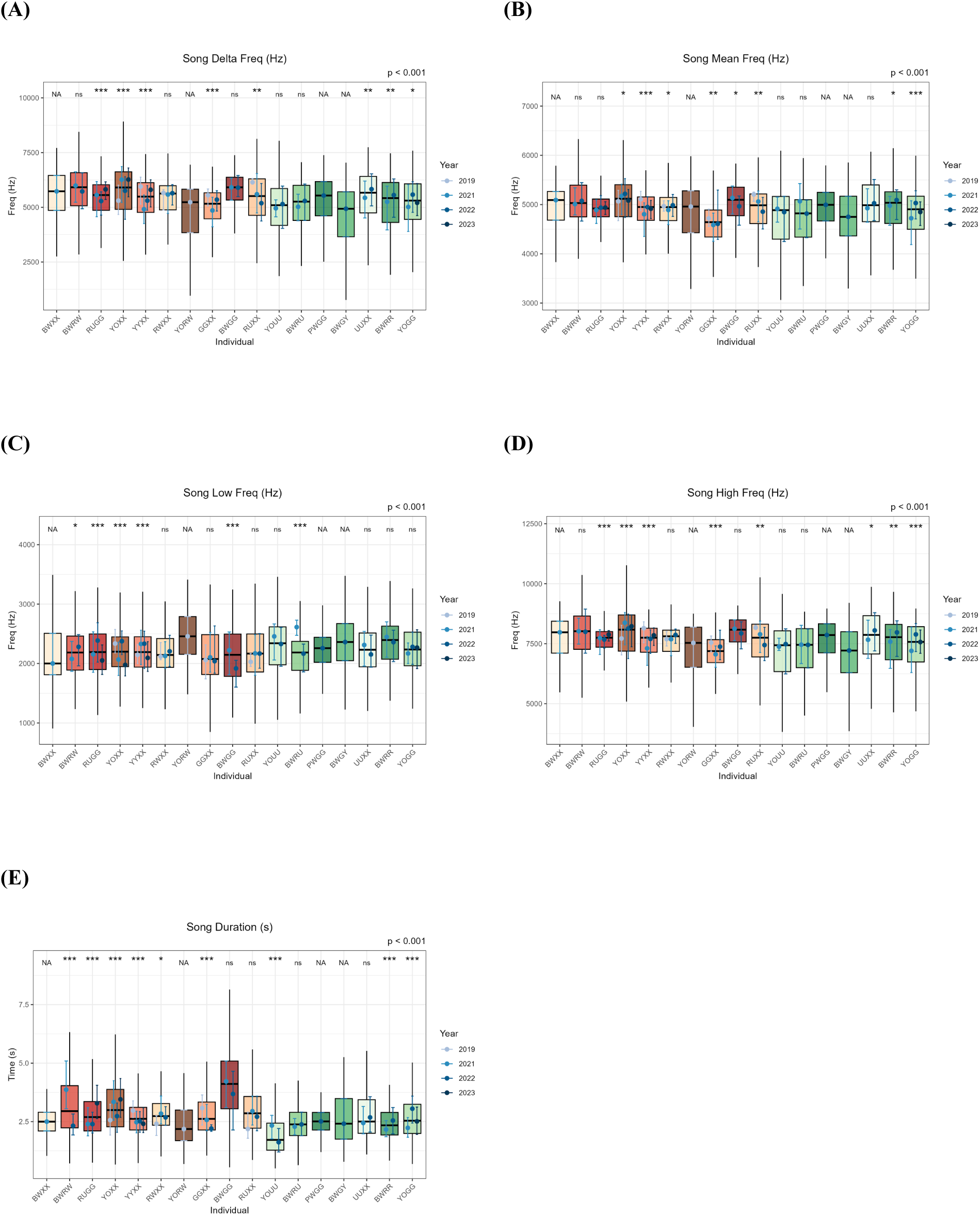
Variation in spectro-temporal parameters across individuals and years. Boxplots show individual-level variation in five song-level acoustic features: (A) Frequency Bandwidth, (B) Mean Frequency, (C) Low Frequency, (D) High Frequency, and (E) Song Duration. Each boxplot represents an individual aggregated across all years (2019–2023), with overlaid colored error bars indicating inter-annual variation within each individual. Asterisks above boxes denote statistical significance of individual identity in linear mixed models (p < 0.05*, ** < 0.01**, *** < 0.001***; “ns” = not significant).

**Figure S5:**
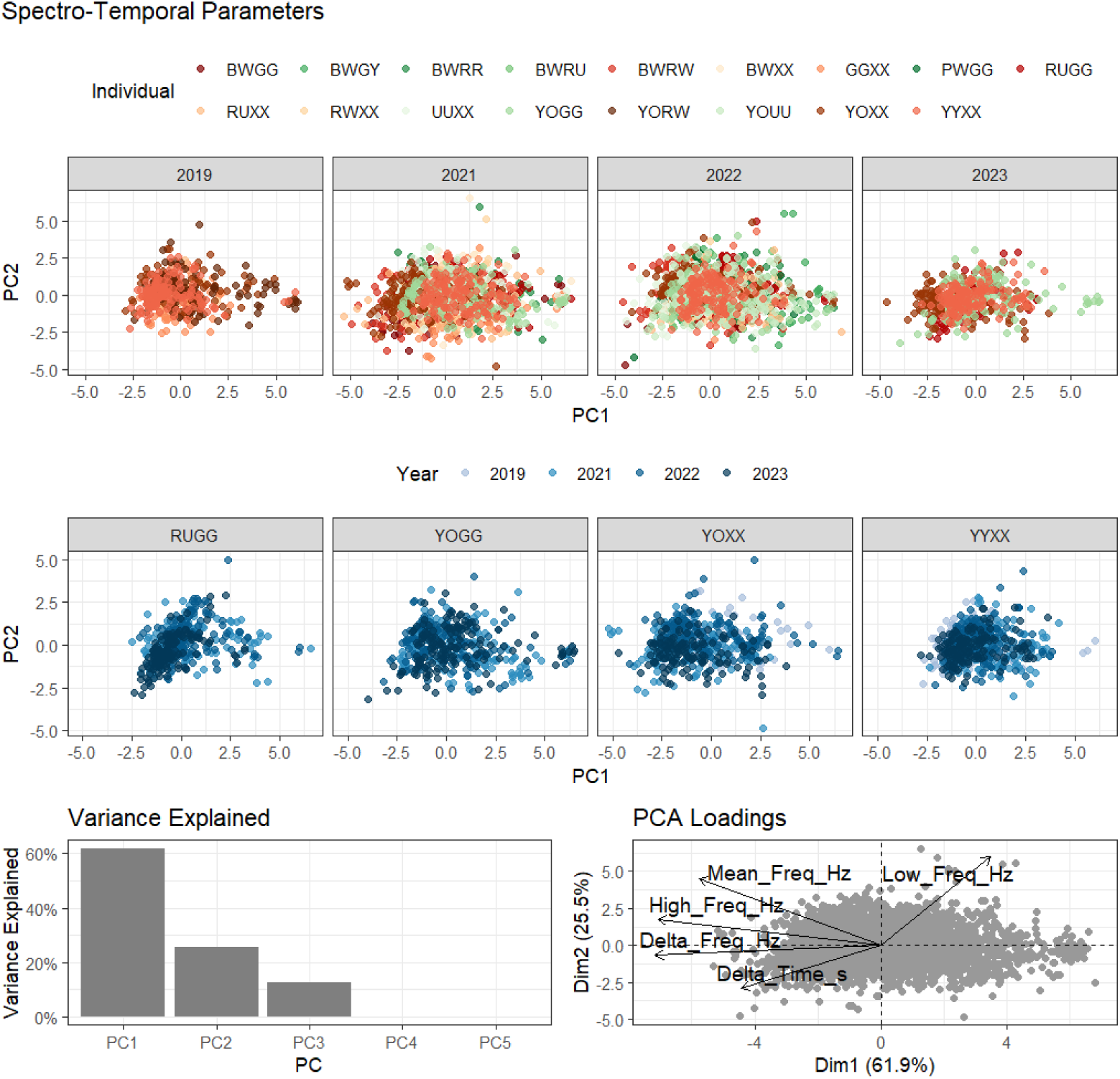
Principal Component Analysis (PCA) of spectro-temporal song parameters. Top panels show PCA scores (PC1 vs. PC2) for all songs, colored by individual identity and faceted by year (2019–2023), illustrating population-wide overlap and inter-annual stability in acoustic space. Middle panels show within-individual PCA scores for four focal males, highlighting individual-specific clustering patterns colored by year. The bottom-left panel shows the variance explained by the top five principal components, with PC1 and PC2 accounting for 62% and 25% of the total variance, respectively. The bottom-right panel shows PCA loadings, indicating that PC1 primarily reflects frequency-based traits, while PC2 captures variation in frequency and temporal spread.

**Figure S6:**
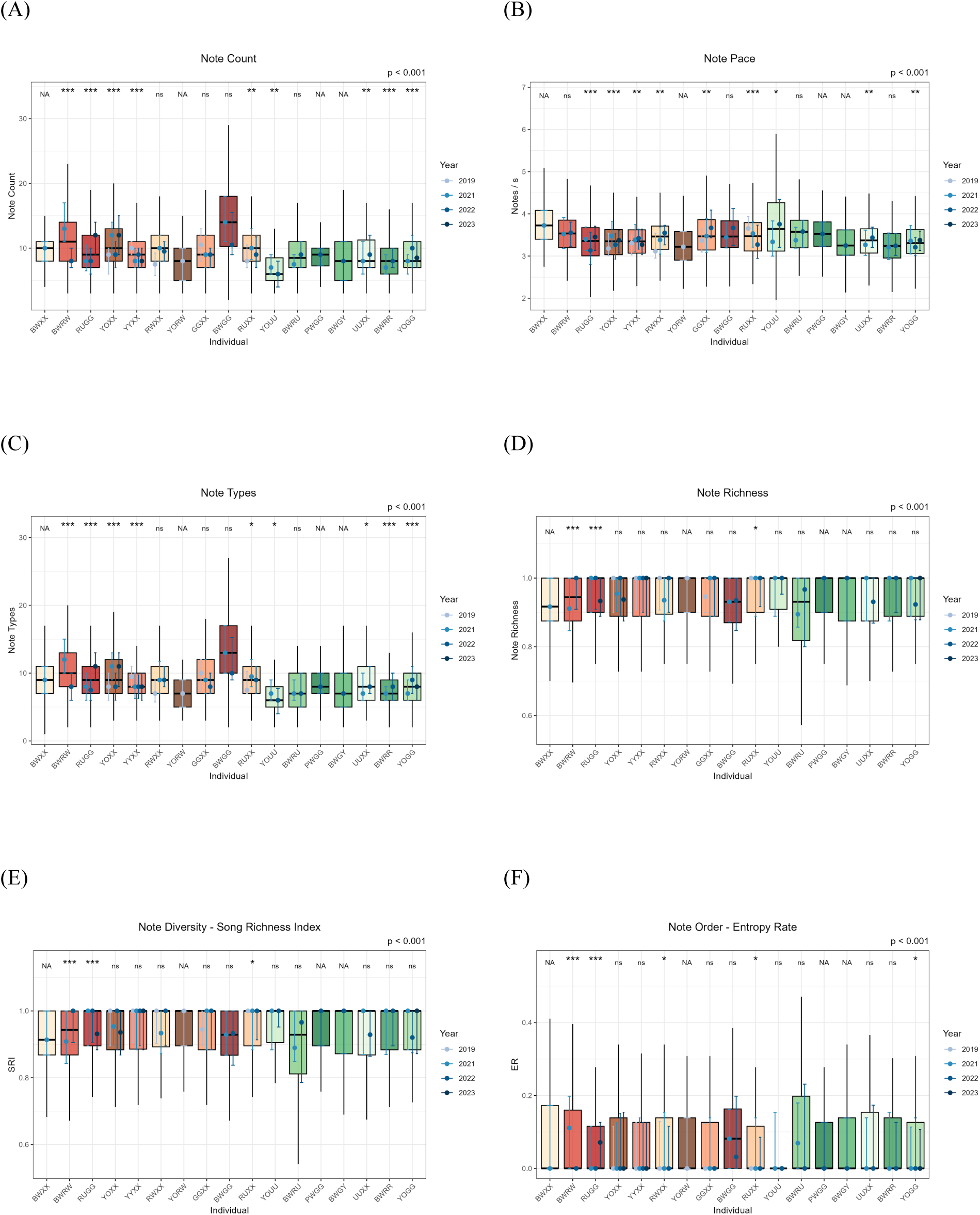
Variation in song complexity metrics across individuals and years. Boxplots show individual-level variation in five song-level acoustic features: (A) Note Count, (B) Note Pace, (C) Note Types, (D) Note Richness, (E) Song Richness Index, and (F) Markov Entropy Rate. Each boxplot represents an individual aggregated across all years (2019–2023), with overlaid colored error bars indicating inter-annual variation within each individual. Asterisks above boxes denote statistical significance of individual identity in linear mixed models (p < 0.05*, ** < 0.01**, *** < 0.001***; “ns” = not significant).

**Figure S7:**
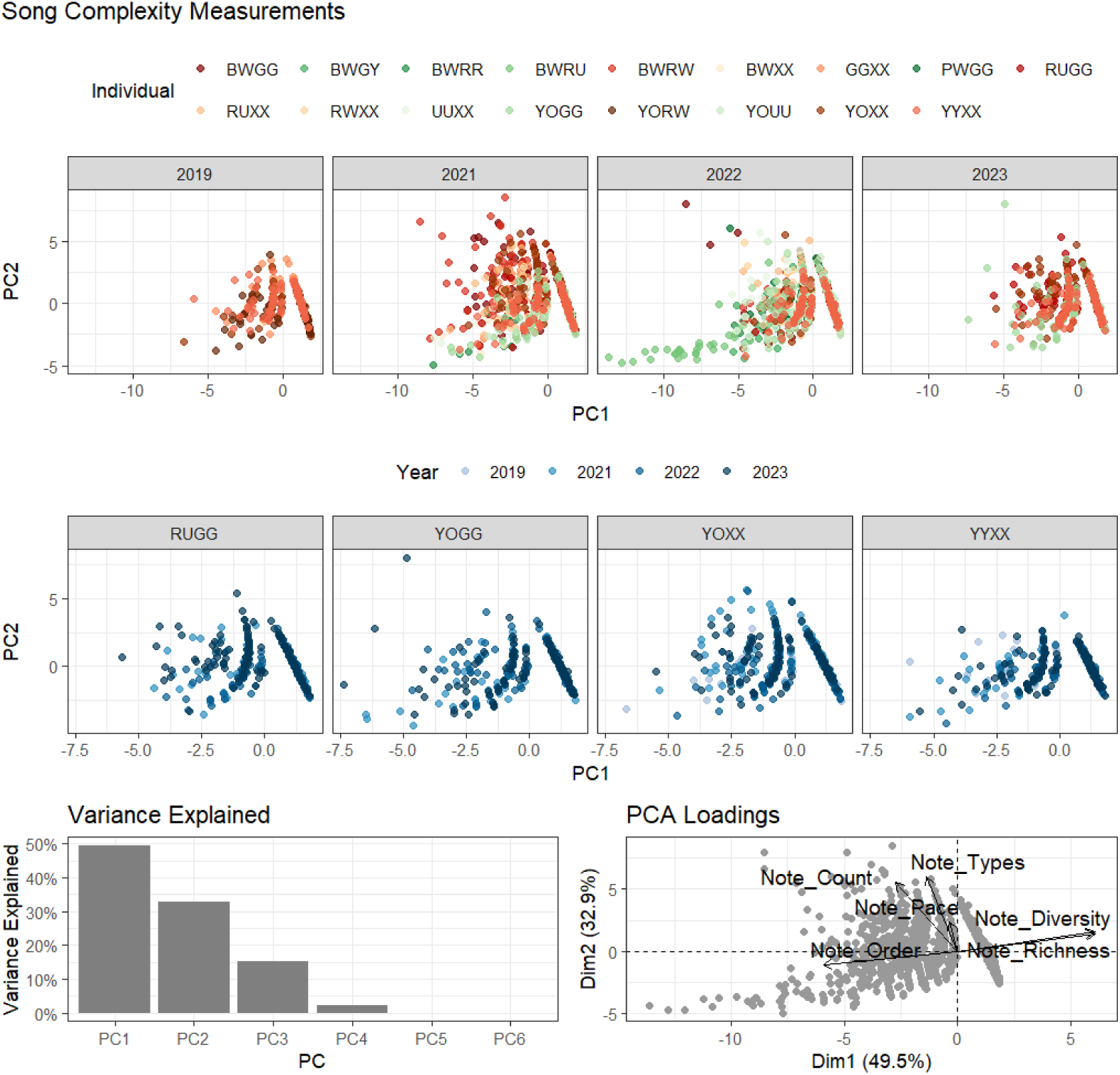
Principal Component Analysis (PCA) of song complexity metrics. Top panels show PCA scores (PC1 vs. PC2) for all songs, colored by individual identity and faceted by year (2019–2023), illustrating population-wide overlap and inter-annual stability in acoustic complexity. Middle panels show within-individual PCA scores for four focal males, highlighting individual-specific clustering patterns colored by year. The bottom-left panel shows the variance explained by the top five principal components, with PC1 and PC2 accounting for 49% and 33% of the total variance, respectively. The bottom-right panel shows PCA loadings, indicating that PC1 primarily reflects frequency-based traits, while PC2 captures variation in frequency and temporal spread.

**Table S1:** A list of all individuals of *S. albiventris* banded during the study period. Band # represents the unique number on the aluminium ring (generally on the right leg), the color-band combination represents the color of rings from top to bottom (generally on the left leg) and the Individual ID represents a unique code used for individual monitoring. * represents the males that were monitored for territorial interactions and acoustic recordings. Individuals were aged based on morphology in the field and sex was determined through molecular sexing. Year of banding is the first time the individual was captured and banded.

| Sr. No. | Band # | Color-band combinations | Individual ID | Sex | Age | Year of Banding |
| --- | --- | --- | --- | --- | --- | --- |
| 1 | A264377 | Pink | KKXX* | Male | Adult | 2018 |
| 2 | A264380 | Green | GGXX* | Male | Adult | 2018 |
| 3 | A264381 | Yellow | YYXX* | Male | Adult | 2018 |
| 4 | A264376 | Red | RRXX* | Unknown | Adult | 2018 |
| 5 | A264383 | White | WWXX* | Unknown | Adult | 2018 |
| 6 | A271204 | Purple-Green | PGXX | Female | Adult | 2019 |
| 7 | A271209 | Yellow-Orange, Purple-Green | YOPG | Female | Adult | 2019 |
| 8 | A271212 | Yellow-Orange, Purple-White | YOPW | Female | Adult | 2019 |
| 9 | A271216 | Yellow-Orange, Green-Yellow | YOGY | Female | Adult | 2019 |
| 10 | A271202 | Black-White | BWXX* | Male | Adult | 2019 |
| 11 | A271203 | Yellow-Orange | YOXX* | Male | Adult | 2019 |
| 12 | A271205 | Blue-Red | URXX* | Male | Adult | 2019 |
| 13 | A271206 | Red-White | RWXX* | Male | Adult | 2019 |
| 14 | A271207 | Yellow-Green | YGXX | Male | Adult | 2019 |
| 15 | A271213 | Yellow-Orange, Red-White | YORW* | Male | Adult | 2019 |
| 16 | A271215 | Yellow-Orange, Red-Blue | YORU | Male | Juvenile | 2019 |
| 17 | A271217 | Black-White, Purple-Green | BWPG | Male | Juvenile | 2019 |
| 18 | A271208 | Yellow-Orange, Black-White | YOBW | Unknown | Adult | 2019 |
| 19 | A271231 | Black-White, Red-White | BWRW* | Male | Adult | 2020 |
| 20 | A269452 | Purple-White, Blue | PWUU | Female | Juvenile | 2021 |
| 21 | A269454 | Red-Blue, White | RUWW | Female | Adult | 2021 |
| 22 | A269456 | Orange-Yellow, Red | OYRR | Female | Adult | 2021 |
| 23 | A269457 | Yellow, Blue-Red | YYUR | Female | Juvenile | 2021 |
| 24 | A269458 | Red-White, Green-Yellow | RWGY | Female | Juvenile | 2021 |
| 25 | A269460 | Purple-Green, Blue | PGUU | Female | Juvenile | 2021 |
| 26 | A269461 | White-Red, White | WRWW | Female | Juvenile | 2021 |
| 27 | A271262 | Black-White, Purple-White | BWPW | Female | Adult | 2021 |
| 28 | A269443 | Red-Blue, Green | RUGG* | Male | Adult | 2021 |
| 29 | A269444 | White-Purple | WPXX* | Male | Adult | 2021 |
| 30 | A269445 | Blue | UUXX* | Male | Adult | 2021 |
| 31 | A269447 | Yellow-Orange, Blue | YOUU* | Male | Adult | 2021 |
| 32 | A269448 | Red-White, Green | RWGG* | Male | Adult | 2021 |
| 33 | A269449 | Yellow-Orange, Green | YOGG* | Male | Adult | 2021 |
| 34 | A269450 | Black-White, Red | BWRR* | Male | Adult | 2021 |
| 35 | A269451 | Red-White, Blue | RWUU* | Male | Juvenile | 2021 |
| 36 | A269453 | Purple-Green, White | PGWW* | Male | Adult | 2021 |
| 37 | A269455 | Purple-White, Green | PWGG* | Male | Adult | 2021 |
| 38 | A269459 | Green-Yellow, White | YGWW | Male | Juvenile | 2021 |
| 39 | A271247 | Black-White, Red-Blue | BWRU* | Male | Adult | 2021 |
| 40 | A271253 | Black-White, Green-Yellow | BWGY* | Male | Adult | 2021 |
| 41 | A271260 | Black-White, Green | BWGG* | Male | Adult | 2021 |
| 42 | A264423 | White, Blue | WWUU* | Male | Adult | 2022 |
| 43 | A264424 | Green-Yellow, Blue-Red | GYUR* | Male | Adult | 2022 |
| 44 | A264425 | White-Black, Yellow | WBYY | Male | Juvenile | 2022 |
| 45 | A264426 | Black-White, Blue | BWUU | Male | Juvenile | 2022 |
| 46 | A264427 | Green, red | GGRR | Female | Adult | 2022 |

**Table S2:**
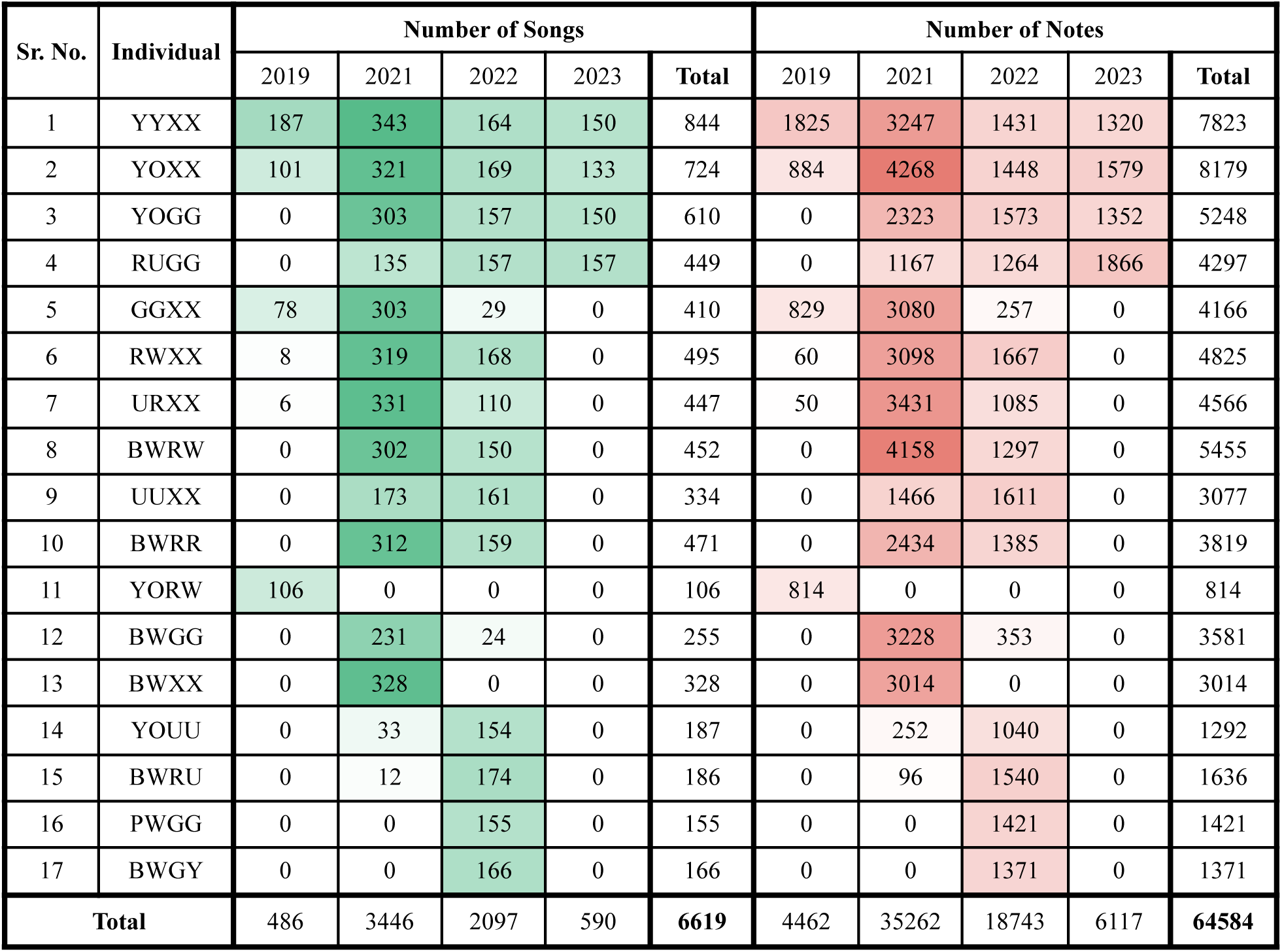
Per year sample size for each color-banded male recorded during the study period. Number of Songs and the total number of notes in the songs annotated per year for each individual. All the songs across individuals and years were used to examine spectro-temporal variation and song complexity. For song syntax analysis, we included individual data in a given year if we had >75 songs and the number of songs per individual per year was capped to 150.

| Sr. No. | Individual | Number of Songs |  |  |  |  | Number of Notes |  |  |  |  |
| --- | --- | --- | --- | --- | --- | --- | --- | --- | --- | --- | --- |
|  |  | 2019 | 2021 | 2022 | 2023 | Total | 2019 | 2021 | 2022 | 2023 | Total |
| 1 | YYXX | 187 | 343 | 164 | 150 | 844 | 1825 | 3247 | 1431 | 1320 | 7823 |
| 2 | YOXX | 101 | 321 | 169 | 133 | 724 | 884 | 4268 | 1448 | 1579 | 8179 |
| 3 | YOGG | 0 | 303 | 157 | 150 | 610 | 0 | 2323 | 1573 | 1352 | 5248 |
| 4 | RUGG | 0 | 135 | 157 | 157 | 449 | 0 | 1167 | 1264 | 1866 | 4297 |
| 5 | GGXX | 78 | 303 | 29 | 0 | 410 | 829 | 3080 | 257 | 0 | 4166 |
| 6 | RWXX | 8 | 319 | 168 | 0 | 495 | 60 | 3098 | 1667 | 0 | 4825 |
| 7 | URXX | 6 | 331 | 110 | 0 | 447 | 50 | 3431 | 1085 | 0 | 4566 |
| 8 | BWRW | 0 | 302 | 150 | 0 | 452 | 0 | 4158 | 1297 | 0 | 5455 |
| 9 | UUXX | 0 | 173 | 161 | 0 | 334 | 0 | 1466 | 1611 | 0 | 3077 |
| 10 | BWRR | 0 | 312 | 159 | 0 | 471 | 0 | 2434 | 1385 | 0 | 3819 |
| 11 | YORW | 106 | 0 | 0 | 0 | 106 | 814 | 0 | 0 | 0 | 814 |
| 12 | BWGG | 0 | 231 | 24 | 0 | 255 | 0 | 3228 | 353 | 0 | 3581 |
| 13 | BWXX | 0 | 328 | 0 | 0 | 328 | 0 | 3014 | 0 | 0 | 3014 |
| 14 | YOUU | 0 | 33 | 154 | 0 | 187 | 0 | 252 | 1040 | 0 | 1292 |
| 15 | BWRU | 0 | 12 | 174 | 0 | 186 | 0 | 96 | 1540 | 0 | 1636 |
| 16 | PWGG | 0 | 0 | 155 | 0 | 155 | 0 | 0 | 1421 | 0 | 1421 |
| 17 | BWGY | 0 | 0 | 166 | 0 | 166 | 0 | 0 | 1371 | 0 | 1371 |
| Total |  | 486 | 3446 | 2097 | 590 | 6619 | 4462 | 35262 | 18743 | 6117 | 64584 |

